# Abstract representations emerge naturally in neural networks trained to perform multiple tasks

**DOI:** 10.1101/2021.10.20.465187

**Authors:** W. Jeffrey Johnston, Stefano Fusi

## Abstract

Humans and other animals demonstrate a remarkable ability to generalize knowledge across distinct contexts and objects during natural behavior. We posit that this ability to generalize arises from a specific representational geometry, that we call abstract and that is referred to as disentangled in machine learning. These abstract representations have been observed in recent neurophysiological studies. However, it is unknown how they emerge. Here, using feedforward neural networks, we demonstrate that the learning of multiple tasks causes abstract representations to emerge, using both supervised and reinforcement learning. We show that these abstract representations enable few-sample learning and reliable generalization on novel tasks. We conclude that abstract representations of sensory and cognitive variables may emerge from the multiple behaviors that animals exhibit in the natural world, and, as a consequence, could be pervasive in high-level brain regions. We also make several specific predictions about which variables will be represented abstractly.

## 1 Introduction

The ability to generalize existing knowledge to novel stimuli or situations is essential to complex, rapid, and accurate behavior. As an example, when shopping for produce, humans make many different decisions about whether or not different pieces of produce are ripe – and, consequently, whether to purchase them. The knowledge we use in the store is often learned from experience with that fruit at home – thus, generalizing across distinct contexts. Further, the knowledge that we apply to a fruit that we buy for the first time might be derived from similar fruits – generalizing, for instance, from an apple to a pear. The determinations themselves are often multi-dimensional and multi-sensory: both firmness and appearance are important for deciding whether an avocado is the right level of ripeness. Yet, at the end of this complex process, we make a binary decision about each piece of fruit: we add it to our cart, or do not – and get feedback later about whether that was the right decision. This produce shopping example is not unique. Humans and other animals exhibit an impressive ability to generalize across contexts and between different objects in many situations.

We hypothesize that this ability to generalize is tied to the geometry of neural representations. Neural representations of sensory and cognitive variables are often highly nonlinear and have high embedding dimension[1–3]. While the nonlinearity of the representations allows flexible learning of new behaviors[2] and provides metabolically efficient and reliable representations[4], high-dimensional representations often do not permit generalization across contexts or stimuli[2, 5]. Alternatively, for a representation with low embedding dimension, a classifier that learns to discriminate between a single pair of stimuli based on one latent variable may generalize to discriminate between other pairs of stimuli that differ in other latent variables. Recent experimental work has shown that low-dimensional, near-linear representations that could support this ability to generalize exist at the apex of the primate ventral visual stream, for faces in inferotemporal cortex[6–8]. Further, experimental work in the hippocampus and prefrontal cortex has shown that representations of the sensory and cognitive features related to a complex cognitive task, also support generalization[5]. We refer to representations of task-relevant sensory and cognitive variables that support generalization – like in these examples and others[9, 10] – as abstract representations.

In the machine learning literature, abstract representations are often referred to as factorized[11] or disentangled[7, 11–14] representations of interpretable stimulus features. Deep learning has been used to produce abstract representations primarily in the form of unsupervised generative models[12, 15, 16] (but see [17]). In this context, abstract representations are desirable because they allow potentially novel examples of existing stimulus classes to be produced by linear interpolation in the abstract representation space (for example, starting at a known exemplar and changing its orientation by moving linearly along a dimension in the abstract representation space that is known to correspond to orientation)[12].

Here, we ask how abstract representations – like those observed in higher brain regions[5, 6] – can be constructed from the nonlinear and high-dimensional representations observed in early sensory areas[3, 18–22]. To study this, we construct high-dimensional and nonlinear representations of latent variables; then, we show that training feedforward neural network models to perform multiple distinct classification tasks on these latent variables induces abstract representations in a wide variety of conditions.

While experimental work on animals performing more than a couple of distinct behavioral tasks remains nearly nonexistent[23], modeling work using recurrent neural networks has shown that the networks often develop representations that can be reused across distinct, but related tasks[24] – though the abstractness of these reusable representations was not measured. Thus, the behavioral constraint of multi-tasking may encourage the learning of abstract representations of stimulus features that are relevant to multiple tasks. To investigate this hypothesis, we train feedforward neural network models to perform multiple distinct tasks on a common stimulus space. Previous work in machine learning has shown that similar multi-tasking networks can achieve lower loss from the same number of samples than networks trained independently on each task[25] (and see [26]), and that they can quickly learn novel, but related, tasks that are introduced after training[27]. Both of these properties are hallmarks of abstract representations – however, to our knowledge, the representational geometry developed by these multi-tasking networks has not been characterized.

We begin by introducing the multi-tasking model and show that it produces fully abstract representations that are surprisingly robust to task heterogeneity and context-dependence. These representations also emerge in the more realistic case in which only a fraction of tasks are closely related to the latent variables, and the remaining larger fraction are not. Next, we characterize how the level of abstraction depends on nonlinear curvature in the classification task boundaries and on different types of inputs, including images. We also show that the multi-tasking model works similarly when trained using reinforcement learning. Finally, we use this framework to make several predictions for how neural representations in the brain will be shaped by behavioral demands. Overall, our work shows that abstract representations – similar to those observed in the brain[5–7, 9] – reliably emerge from learning to multi-task in multi-dimensional environments. Together, our results indicate that abstract representations in the brain may be a consequence of – as well as a boon to[28] – complex behavior.

## 2 Results

### 2.1 Abstract representations allow knowledge to be generalized across contexts

The knowledge of latent structure that is present in the sensory world can enable generalization. For example, the appearance of different kinds of berries can be described by two continuous latent variables: color and shape. As an example, berries that have a similar shape are likely to also have similar texture when eaten, regardless of their color (fig. 1a, top); further, berries that are red may taste more similar to each other, despite differences in shape, than they do to berries that are blue (fig. 1a, bottom). Learning and taking advantage of this structure in the sensory world is important for animals that need to quickly react to novel stimuli using information from previously experienced stimuli.

**Figure 1:**
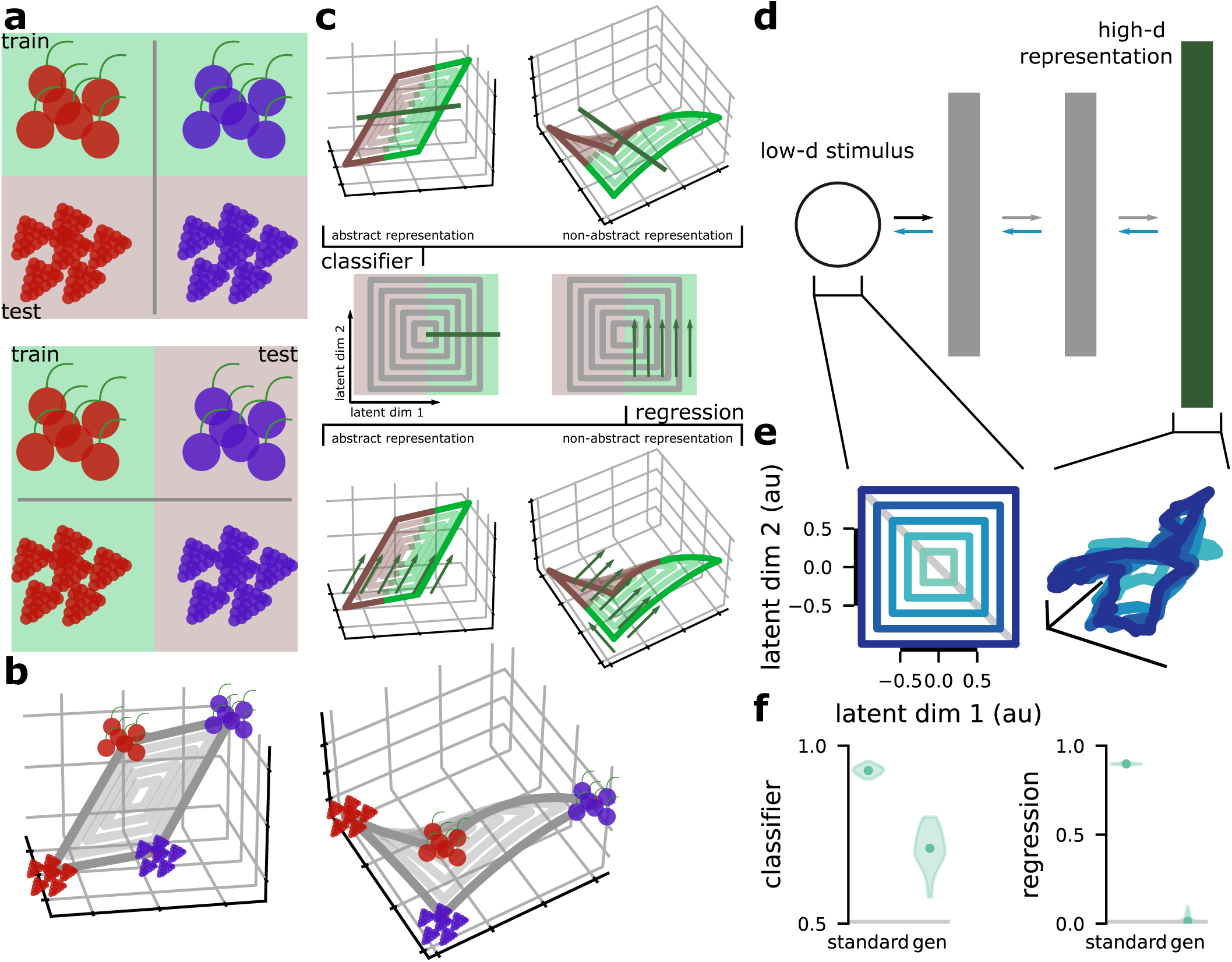
The abstraction metrics and input representations. **a** Illustration of the classification tasks. (top) A classification learned between red and blue berries of one shape should generalize to other shapes. (bottom) A classification between red berries of two different shapes should generalize to blue berries of different shapes. **b** Examples of linear, abstract (left) and nonlinear, non-abstract (right) representations of the four example berries. **c** Illustration of our two abstraction metrics. For a *D* = 2-dimensional latent variable (middle), we split the latent variable distribution into two regions: one used for training (green, left) and one used for testing (red, right). In the classifier generalization metric, we train a linear classifier to perform a binary classification of the latent variables using samples from the green region and test that classifier on samples from the red region (left). The abstract representation (top left) has good classifier generalization performance; the non-abstract representation (top right) has poor classifier generalization performance. The regression generalization metric is similar, but uses a linear regression model (right). The abstract representation (bottom left) has good regression generalization performance; the non-abstract representation (bottom right) has poor regression generalization performance. **d** We use a feedforward network to produce high-dimensional representations from our *D*-dimensional latent variables. The network is trained to maximize the dimensionality of the representation layer (green), while also retaining the ability to reconstruct the original latent variables through an autoencoder (blue lines). **e** We visualize the structure of the latent variable representation by plotting the first three principal components (left: original structure; right: representation structure). **f** We use both the classifier (left) and regression (right) generalization metrics to quantify that level of abstraction. In each plot, the left point is for the metric trained and tested on the whole space, the right point is trained on one half of the space and tested on the other half. The grey line is chance.

We refer to neural representations that reflect this latent structure as abstract. In the example above, one form of an abstract representation of these latent variables is a linear representation of them in neural population activity, such a representation would have low-dimensional, rectangular structure in neural population space (fig. 1b, left); a non-abstract representation of these latent variables would have a higher-dimensional distorted structure, such as one created by neurons that each respond only to particular conjunctions of color and shape (fig. 1b, right). The abstract representation has the desirable quality that, if we learned a neural readout that classifies blue berries from red berries using berries with only one shape (e.g., the two bottom berries in fig. 1b, left), then we would not need to modify this classifier to apply it to berries of a different shape (e.g., the two top berries in fig. 1b, left); while the same classification can be learned for the non-abstract representation, it will not generalize (compare the two berries to the left and to the right in fig. 1b, right).

We quantify abstraction using two metrics, both of which are related to a metric used to quantify abstraction in previous experimental work[5]. The first tests how well a classifier that is trained on one half of the stimulus space generalizes to the other half of the stimulus space (fig. 1c, top). High classifier generalization performance has been observed for sensory and cognitive features in neural data recorded from the hippocampus and prefrontal cortex[5]. The second tests how well a linear regression model that is trained on one half of the stimulus space generalized to the other half of the stimulus space (fig. 1c, bottom; similar metrics are often used in the machine learning literature[12, 29]). The classifier generalization metric requires that the coarse structure of the representations be abstract, but is less sensitive to small deviations. The regression generalization metric is much stricter, and is sensitive to even small deviations from a representation that follows the underlying latent variable structure. In some cases, we also compare these metrics of out-of-distribution generalization to standard cross-validated performance on the whole latent variable space. Intuitively, the standard cross-validated performance of both metrics serves as a best case for their out-of-distribution generalization performance (i.e., the case where what is learned from only half the representation space is just as informative about the global representation structure as what would be learned from the whole representation space). In a perfectly abstract representation, the standard and out-of-distribution generalization performances would be equal to each other. Just as similar metrics were used to quantify the abstractness of neural representations recorded experimentally, we use the classification and regression generalization performance to quantify the abstractness of the representations developed by the multi-tasking model.

### 2.2 Understanding the learning dynamics that produce abstract representations

First, we develop a model to construct non-abstract representations of known *D*-dimensional latent variables which we refer to as the standard input (here, the latent variables are given a standard normal distribution – though the results are similar for a uniform distribution, see *A sensitivity analysis of the multi-tasking model and βVAE* in *Supplement*). These non-abstract representations will be used as the input to the multi-tasking model. To produce the non-abstract standard input, we train a feedforward neural network with an autoencoder to satisfy two objectives: First, to maximize the dimensionality of activity in the representation layer and, second, to reconstruct the original stimulus using only the representation. That is, we want a high dimensional representation that still preserves all of the information about the input. This transformation produces a distorted and tangled representation of the latent variables (fig. 1d) and, for a 5-dimensional latent variable, produces a representation layer with a dimensionality of approximately 200 (see *Participation ratio-maximized representations* in *Methods* for more detail). We visualize this transformation by constructing concentric squares in the latent variable space (fig. 1e, left) and then visualizing the representation of these squares produced by the network (fig. 1e, right). Preserved concentric square structure suggests abstraction, while a lack of that structure suggests a lack of abstraction. The distorted representation of the latent variables produced here significantly decreases abstraction, as measured by both classifier (fig. 1f, left) and regression (fig. 1f, right) generalization metrics.

To recover abstract structure from non-abstract representations, we introduce the multi-tasking model (fig. 2a). The multi-tasking model is a multilayer feedforward neural network model that is trained to perform *P* different binary classification tasks (see *The multi-tasking model* in *Methods* for details). These tasks are analogous to the tasks that animals perform, as described above. For instance, if an animal eats a berry, the animal later receives information about whether that berry was edible or poisonous. If we assume that the edibility of a berry is represented by one of our *D* latent variables, then, in the multi-tasking model, this classification task corresponds to the model being trained to produce one output when the latent variable is positive and another output when the latent variable is negative. In the full model, each classification task is chosen to be a random hyperplane in the full *D*-dimensional latent variable space (i.e., each task depends on multiple latent variables). In all of our analyses, we focus on the representations of the stimuli that are developed in the layer preceding the task output layer, which we refer to as the representation layer.

**Figure 2:**
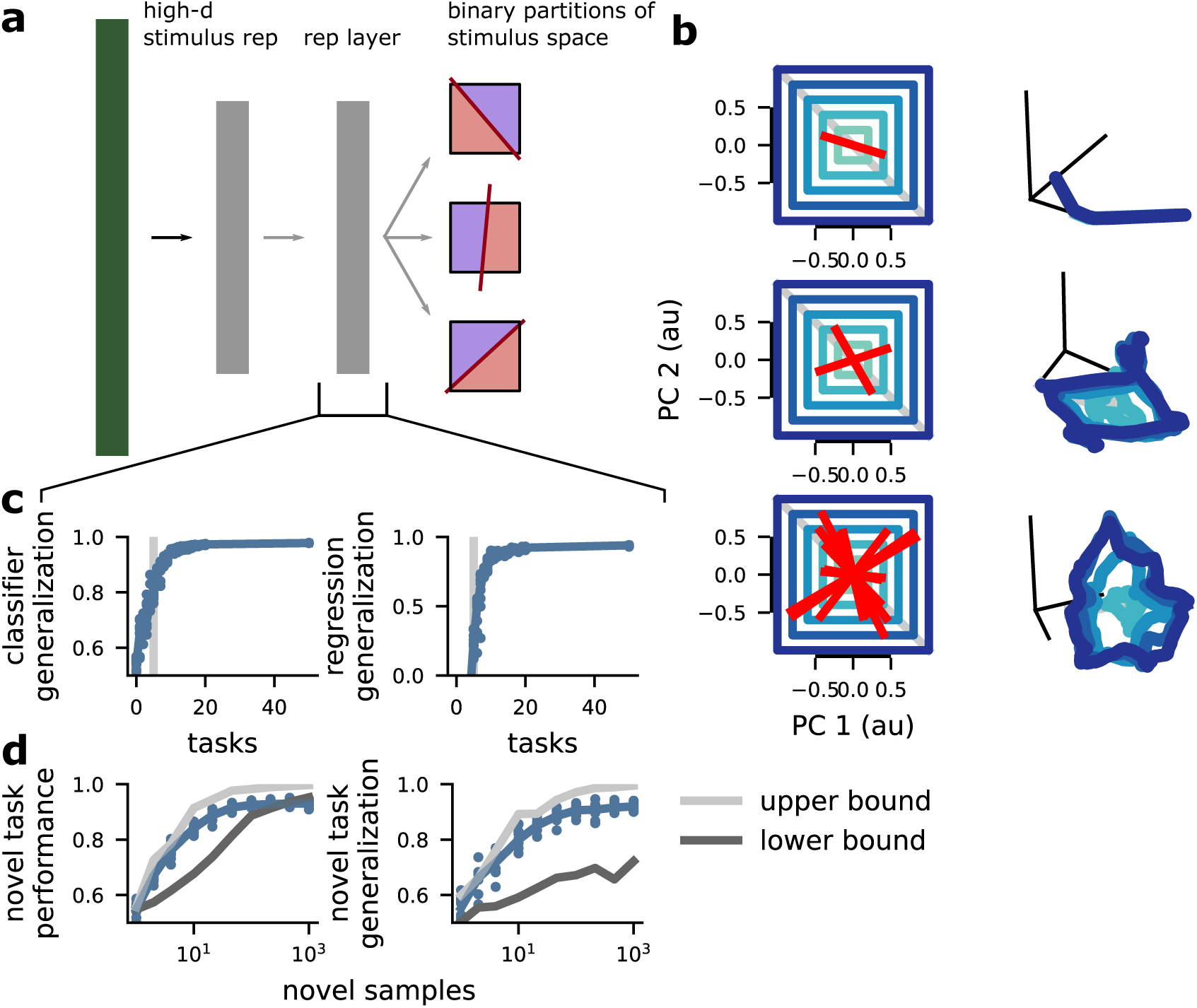
The emergence of abstraction from classification task learning. **a** Schematic of the multi-tasking model. It receives an entangled stimulus representation (as shown in fig. 1e, left) and learns to perform *P* binary classifications of the latent variables. We study the representations that this induces in the layer prior to the output: the representation layer. **b** Examples of multi-tasking model representations for different numbers of classification tasks. We show a schematic of an idealized fully abstract representation (left) alongside the representation developed by the network (right). (top) When the model learns one task (left, red line), the representation (right) collapses into one dimension. (middle) When the model learns two tasks (left, red lines), it recovers more of the stimulus structure (right). (bottom) When the model learns more tasks than stimulus dimensions (here, stimulus dimension is five and eight tasks are learned), the model can produce a highly abstract representation of the original stimuli. **c** The classifier (left) and regression (right) metrics applied to model representations with different numbers of tasks. **d** The standard (left) and generalization (right) performance of a classifier trained to perform a novel task with limited samples using the representations from an multi-tasking model trained with *P* = 10 tasks as input. The lower and upper bound are the standard or generalization performance of a classifier trained on the input representations (lower) and directly on the latent variables (upper). Note that the multi-tasking model performance is close to that of training directly on the latent variables in all cases.

The multi-tasking model is trained to simultaneously produce output for *P* different random tasks. Importantly, the standard input used in this section already has high classification performance for random hyperplane tasks on the latent variables (fig. 1f, left), due to its high-dimensionality[1]. So, one possibility is that the representation layer in the multi-tasking model has the same, non-abstract structure. Here, we show that the feedback onto the representation layer increases the strength of the representation of an approximately min(*P, D*)-dimensional component of the activity. In particular, for a simplified multi-tasking model with a linear output layer, the loss has the form,

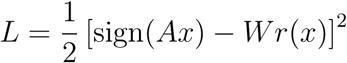

where *A* is a *P × D* matrix of randomly selected partitions (and it is assumed to be full rank). This loss is minimized by making *r*(*x*) a linear transform of sign(*Ax*). In backpropagation, this is achieved by increasing the strength of a component of *r*(*x*) that has the same dimensionality as sign(*Ax*) (see *The dimensionality of representations in the multi-tasking model* in *Methods*). So, we show that,

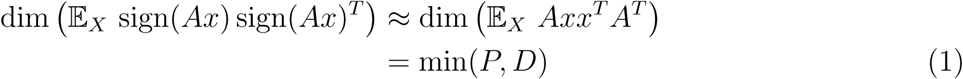

and the approximation becomes closer as *D* becomes larger. This means that, given application of backpropagation, the representation layer will be dominated by a min(*P, D*)-dimensional representation of the latent variables. Since this representation must also be able to satisfy the *P* tasks, it will at least have high classifier generalization performance and may even have high regression generalization performance. While the multi-tasking model used in the rest of the paper has a sigmoid output nonlinearity, the intuition developed in this simplified case still applies.

### 2.3 Learning multiple classification tasks leads to abstract representations

We show that feedforward multilayer neural networks, when trained to perform *P ≥ D* classification tasks, develop fully abstract representations of the *D* latent variables. First, we visualize how the representations developed by our model compare to the abstract latent variables. In particular, we again generate concentric squares in latent variable space and show their idealized abstract representation (fig. 2b, left) alongside the representations actually developed by the model (fig. 2b, right). For only a single task, the representations in the model collapse along a single dimension, which corresponds to performance of that task (fig. 2b, top). While this representation is not abstract, it does mirror distortions in sensory representations that are often observed when animals are overtrained on single tasks[30, 31]. However, when we include a second task in the training procedure, abstract representations begin to emerge (fig. 2b, middle). In particular, the representation layer is dominated by a two-dimensional abstract representation of a linear combination of two of the latent variables. From our theory – and confirmed by these simulations – we know that when *P < D*, then the dimensionality of this dominating component in the representation layer will be approximately *P*. Next, we demonstrate that this abstract structure becomes more complete as the number of tasks included in training is increased. For *P* = 8 and *D* = 5, the visualization suggests that the representation has become fairly abstract (fig. 2b, bottom).

Second, we quantify how the level of abstraction developed in the representation layer depends on the number of classification tasks used to train the model (fig. 2c). For each number of classification tasks, we train 10 multi-tasking models to characterize how the metrics depend on random initial conditions. As the number of classification tasks *P* exceeds the dimensionality of the latent variables *D*, both the classification and regression generalization metrics saturate to near their maximum possible values (classifier generalization metric: exceeds 90 % correct with 8 tasks; regression generalization metric: exceeds *r*^2^ = .8 with 9 tasks; fig. 2c, right of the grey line). Saturation of the classifier generalization metric indicates that the broad organization of the latent variables is perfectly preserved (but detailed information may have been lost); while saturation of the regression generalization metric indicates that even the magnitude information that the multi-tasking model did not receive supervised information about is preserved and represented in a fully abstract format. Importantly, both the training and testing set split and the classification boundary for the classifier generalization metric are randomly selected – they are not the same as classification tasks used in training.

Third, we also show that the multi-tasking model reduces the number of samples required to both learn and generalize novel tasks. For an multi-tasking model trained to perform *P* = 10 tasks with *D* = 5 latent variables, we show how the performance of a novel classification task depends on the number of novel samples, and compare this performance to a lower bound, from when the task is learned from the standard input representation, and an upper bound, from when the task is learned directly from the latent variables. The performance of the multi-tasking model nearly saturates this upper bound (fig. 2d, left). Next, we perform a similar novel task analysis, but where the novel task is learned from one half of the stimulus space and is tested in the other half – just like our generalization analysis above (and schematized in fig. 1c, top). We compare to the same lower and upper bound as before and show that, again, the multi-tasking model representation nearly saturates the upper bound (fig. 2d, right). Thus, not only does the multi-tasking model produce representations with good generalization properties, it also produces representations that lend themselves to the rapid (i.e., few sample) learning of novel tasks.

Finally, we test how robust this finding is to changes to the classification tasks themselves. In particular, we show that our finding holds for three manipulations to the task structure. First, we show that unbalanced tasks (e.g., a more or less stringent criteria for judging the ripeness of a fruit – so either many more of the fruit are considered ripe than spoilt or vice versa; fig. S2a, top left; see *Unbalanced task partitions* in *Methods* for more details) have a negligible effect on the emergence of abstract representations (classifier generalization metric: exceeds 90 % correct with 9 tasks, regression generalization metric: exceeds *r*^2^ = .8 with 9 tasks; fig. S2b). Second, we show that contextual tasks (e.g., determining the ripeness of different fruits that occupy only a fraction of latent variable space; fig. S2a, top right; see *Contextual task partitions* in *Methods* for more details) produce a moderate increase in the number of tasks required to learn abstract representations (classifier generalization metric: exceeds 90 % correct with 14 tasks, regression generalization metric: exceeds *r*^2^ = .8 with 14 tasks; fig. S2b). Third, we show that using training examples with information from only a single task (e.g., getting only a single data point on each trip to the store; fig. S2a, bottom, see *Partial information task partitions* in *Methods* for more details) also only moderately increase the number of tasks necessary to produce abstract representations (classifier generalization metric: exceeds 90 % correct with 11 tasks, regression generalization metric: exceeds *r*^2^ = .8 with 14 tasks; fig. S2b).

Together, these results indicate the the multi-tasking model reliably produces abstract representations even given substantial heterogeneity in the amount of information per stimulus example and the form of that information relative to the latent variables. Further, these results are also robust to variation in architecture: Changing the width, depth, and several other parameters of the multitasking model have only minor effects on classification and regression generalization performance (see *A sensitivity analysis of the multi-tasking model and βVAE* in *Supplement*).

### 2.4 Abstract representations only emerge when task-relevant

Here, we show that abstract representations for the latent variables do not emerge when the multitasking model is trained to perform random, highly nonlinear tasks. We argue that this follows what would be expected in the natural world: latent variables are learned as a way to solve multiple related tasks and to generalize knowledge from one task to another, rather than for their own sake. Then, we show that that abstract representations are recovered when the multi-tasking model learns a combination of aligned and unaligned tasks.

First, we construct grid classification tasks, in which the latent variable space is divided into grid chambers, where each chamber has a roughly equal probability of being sampled (fig. 3a, red lines). Then, we randomly assign each of the grid chambers to one of two categories (fig. 3a, coloring; see *Grid classification tasks* in *Methods* for more details). In this case, there is nothing in the design of the multi-tasking model that privileges a representation of the original latent variables, since they are no longer useful for learning to perform the multiple grid classification tasks. Consequently, the multi-tasking model does not recover a representation of the original latent variables (fig. 3b).

**Figure 3:**
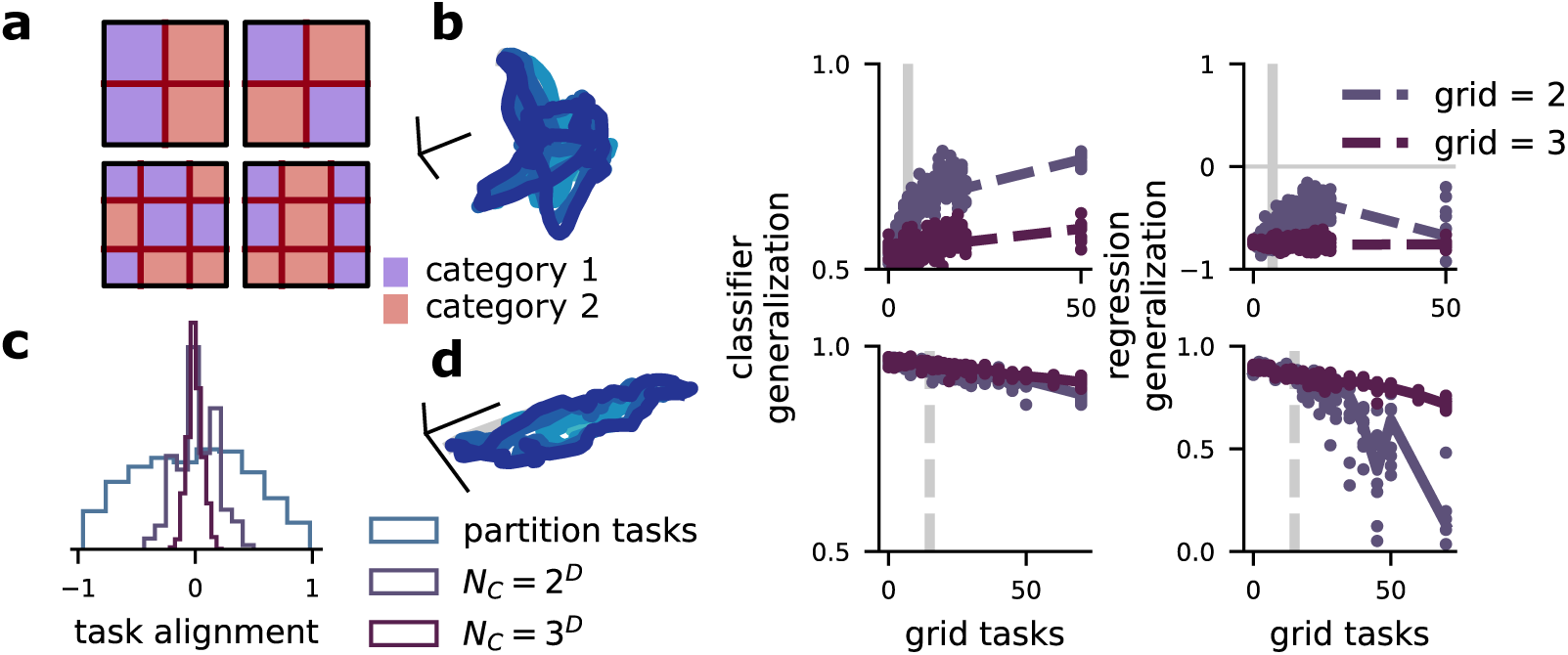
Abstract representations emerge for heterogeneous tasks, and in spite of high-dimensional grid tasks. **a** Schematic of the new grid tasks. They are defined by *n*, the number of regions along each dimension (top: *n* = 2; bottom: *n* = 3), and the dimensionality of the latent variables, *D*. There are *n*^*D*^ total grid chambers, which are randomly assigned to category 1 (red) or category 2 (blue). Some grid tasks are aligned with the latent variables by chance (as in top left), but this fraction is small for even moderate *D*. **b** A multi-tasking model trained only on grid tasks. (left) Visualization of the representation. (middle) Classifier generalization performance. (right) Regression generalization performance. **c** The alignment (cosine similarity) between between randomly chosen tasks for latent variable aligned classification tasks, *n* = 2 and *D* = 5 grid tasks, and *n* = 3 and *D* = 5 grid tasks. **d** As **b**, but for a multi-tasking model trained with *P* = 15 latent variable aligned classification tasks and a variable number of grid tasks.

To make this intuition about the grid tasks more explicit, we show that – in contrast to the latent variable-aligned tasks that we have been using so far – the outcomes from a particular grid task are likely to be only weakly correlated with the outcomes from a different, randomly chosen grid task (fig. 3c). Thus, rather than having a *D*-dimensional structure for *P ≫ D* tasks, the grid tasks will have a roughly *P*-dimensional structure for *P* tasks. As expected, the multi-tasking model fails to learn a strongly abstract representation of the original latent variables, and the representation becomes less abstract as the grid tasks become higher dimensional (i.e., when the grid has more chambers; fig. 3b, middle and right, blue and purple lines).

Next, we examine the representations learned by the multi-tasking model when it must perform a mixture of latent variable-aligned and grid classification tasks. This situation is also chosen to mimic the natural world, as a set of latent variables may be relevant to some behaviors (the latent variable-aligned classification tasks), but an animal may need to perform additional behaviors on the same set of stimuli that do not follow the latent variable structure (the grid classification tasks). Here, we train the multi-tasking model to perform a fixed number of latent variable-aligned tasks, which are sufficient to develop abstract structure in isolation (here, 15 tasks). However, at the same time, the model is also being trained to perform various numbers of grid tasks. While increasing the number of grid tasks does moderately decrease the abstractness of the developed representation (fig. 3d, middle and right), the multi-tasking model retains strongly abstract representations even while performing more than 45 grid tasks – 3 times as many as the number of latent variable-aligned tasks.

Intuitively, this occurs because the latent variable-aligned tasks are correlated with each other and follow the structure of the *D*-dimensional latent variable space, while each of the grid tasks has low correlation with any other grid task (fig. 3b). Thus, a shared representation structure is developed to solve all the latent variable-aligned tasks essentially at once (corresponding to significant fraction of the variance in the target function), while a smaller nonlinear component is added on to solve each of the grid tasks relatively independently. Interestingly, the combination of abstract structure with nonlinear distortion developed by the multi-tasking model here has also been observed in the brain and other kinds of feedforward neural networks (though learning tasks analogous to our grid tasks was not necessary for it to emerge)[5]. We believe that this compromise between strict abstractness to allow for generalization and nonlinear distortion to allow for flexible learning of random tasks[1, 2] is fundamental to the neural code.

### 2.5 The multi-tasking model learns abstract structure from tasks with nonlinear curvature

While we have shown that the multi-tasking model learns abstract structure from several different manipulations of linear tasks (fig. S2a, b) and fails to learn abstract structure from highly nonlinear tasks, for which the latent variables themselves are no longer relevant (fig. 3a, b), these two examples represent relatively extreme cases. Here, we show that the multi-tasking model still produces abstract representations in many cases in between these two extremes, when it is trained on tasks with different levels of nonlinear curvature (fig. 4b, d). To produce these tasks, we generate random Gaussian processes with radial basis function kernels of a particular length scale (fig. 4b, right), then use it to produce two distinct categories by binarizing the output (where outputs *>* 0 are in one category and *≤* 0 are the other; fig. 4b, left, and see *Random Gaussian process tasks* in *Methods* for more details).

**Figure 4:**
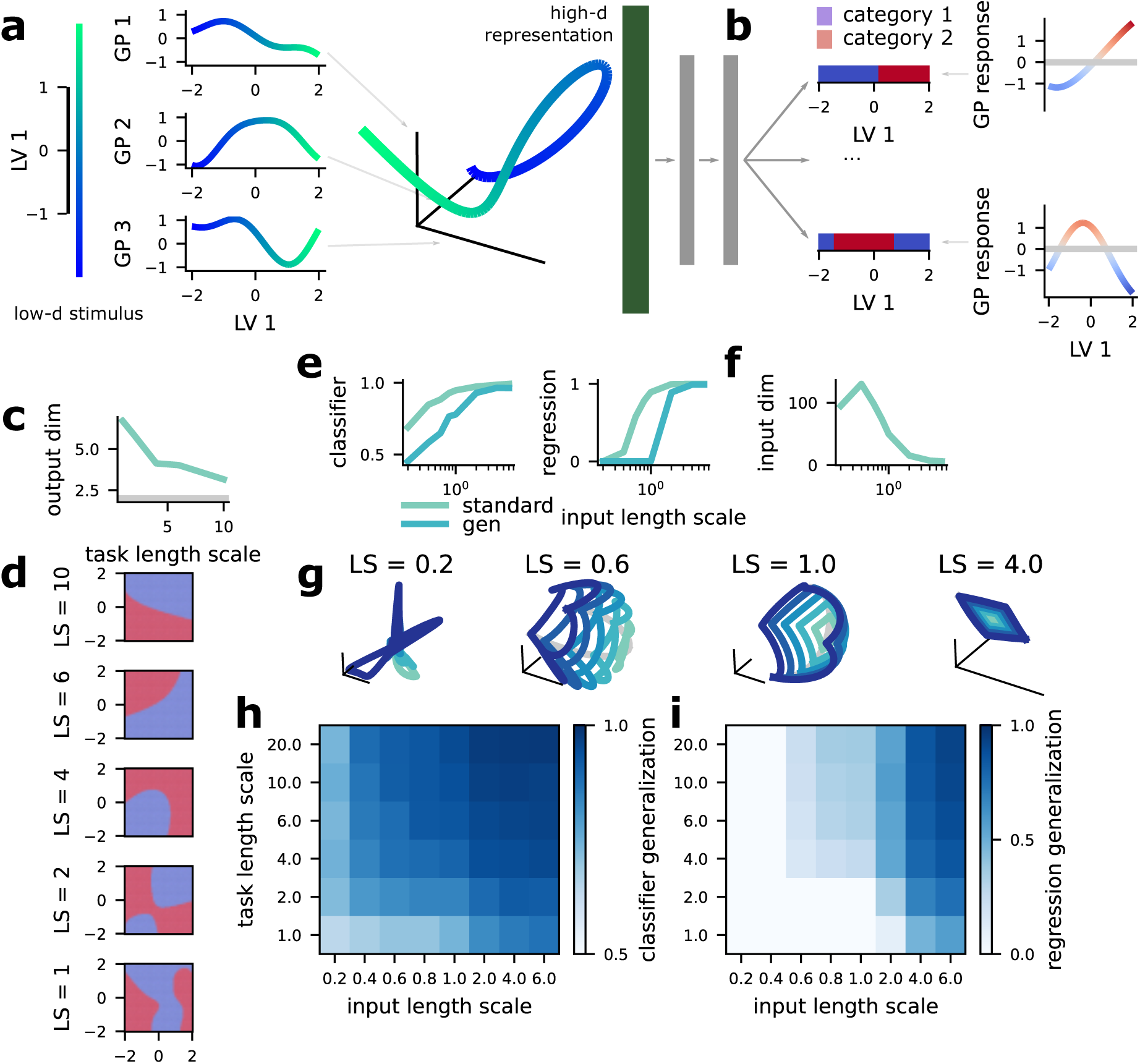
The multi-tasking model learns abstract structure from both random Gaussian process inputs and output tasks. **a** Schematic of the creation of a random Gaussian process input. (left) Example traversal of one latent variable dimension (*D* = 1). (middle) Random Gaussian processes with length scale = 1 learned for the single latent variable shown on the left. (right) The response produced by these three random Gaussian process for the example traversal. The full random Gaussian process input has 500 random Gaussian process dimensions and is used as input to the multi-tasking model. **b** Schematic of the creation of two random Gaussian process tasks for the *D* = 1-dimensional latent variable shown in **a**. (left) The two tasks binary classification tasks. (right) The underlying random Gaussian process for each of the two tasks on the left. **c** The dimensionality (participation ratio) of the binary output patterns required by task collections of different length scales. **d** Examples of random Gaussian process tasks for a variety of length scales. The multi-tasking model is trained to perform these tasks, as schematized in **b. e** Classifier (left) and regression (right) generalization performance for random Gaussian process inputs of different length scales. Note that for long length scales, the input is already abstract. **f** The dimensionality (participation ratio) of the random Gaussian process for different length scales. Note that it is always less than the dimensionality of 200 achieved by the standard input. **g** Visualization of the input structure for random Gaussian process inputs of different length scales. **h** Classifier generalization performance of an multi-tasking model trained to perform *P* = 15 classification tasks with *D* = 5-dimensional latent variables, shown for different conjunctions of task length scale (changing along the y-axis) and input length scale (changing along the x-axis). **i** Regression generalization performance shown as in **h**.

Following this procedure, we produce tasks with a variety of length scales (fig. 4d, length scale decreases from top to bottom). Tasks with lower length scales will have more curved boundaries and multiple distinct category regions, similar to the grid tasks (fig. 4d, bottom); tasks with higher length scales will tend to have less curved boundaries – and large length scales (e.g., *>* 15) will approximate the linear tasks from before. We quantify the nonlinearity of these tasks by computing how the dimensionality of the required output depends on task length scale for *P* = 15 classification tasks on *D* = 2 latent variables (fig. 4c). As discussed above, the nonlinear grid tasks have an output dimensionality that approaches the number of tasks, while the linear tasks used above have an output dimensionality that is only slightly higher than the number of latent variables. We show that the random Gaussian process tasks have a required output dimensionality that lies between these extremes, and that decreases with increased length scale (fig. 4c). This suggests that the multi-tasking model will learn abstract representations for moderate levels of curvature.

We show that an multi-tasking model trained to perform *P* = 15 classification tasks produces representations with above-chance classifier generalization performance for all task length scales that we investigated (fig. 4h, left, middle columns). Further, the multi-tasking model produces representations with above-chance regression generalization performance for many different task length scales as well (fig. 4h, right, middle columns), though it is less consistent than classifier generalization performance. Thus, the multi-tasking model produces partially abstract representations even from highly curved and sometimes multi-region task boundaries, and produces fully abstract representations for curved task boundaries. So, while the multi-tasking model does not produce abstract representations in the extreme case of the highly nonlinear grid tasks, it does produce abstract representations for many intermediate task structures (shown both here and above).

### 2.6 The multi-tasking model learns abstract structure from random Gaussian process inputs

To explore how the learning of abstract structure depends on the kind of input that we supply to the multi-tasking model, we use a new kind of nonlinear input: random Gaussian processes with radial basis function kernels of different length scales (fig. 4a, g). To illustrate this approach, we begin with a *D* = 1 normally distributed latent variable (fig. 4a, left), then generate random Gaussian process functions that map this variable to a scalar output (three functions in fig. 4a, center). The scalar outputs of many random Gaussian process functions are then used as input to the multi-tasking model (fig. 4a, right; and see *Random Gaussian process inputs* in *Methods* for more details). Where we show results from these random Gaussian process inputs, we use *D* = 5 rather than the *D* = 1 used in the example.

The length scale of the kernel controls how nonlinear and tangled the representation is (fig. 4g). At one extreme, a random Gaussian process with a low length scale (e.g., *<* 1) will have low classification and regression generalization performance; at the other extreme, a random Gaussian process with high length scale (e.g., *>* 5) will have high classification and regression generalization performance (fig. 4e). This roughly follows the dimensionality of the input after transformation by the random Gaussian process (fig. 4f). We show that the multi-tasking model achieves high classifier generalization performance for random Gaussian process inputs with many different length scales, including low length scales where the input has no existing abstract structure (fig. 4h, top row). We also show that the multi-tasking model achieves moderate regression generalization performance for many different length scales as well, though regression generalization performance remains at chance for the shortest length scales that we investigated (fig. 4i, top row).

The random Gaussian process input differs from our previous input type in that, for low length scales, a linear decoder cannot reliably learn random categorical partitions (as is the case for the standard input, see fig. 1f). The random Gaussian process representations also have significantly lower participation ratios than those produced by the standard input. We can see that the random Gaussian process input tends to fold back on itself for low length scales (fig. 4a), exhibiting a periodic structure that we do not observe in the standard input (not shown). This increased periodic structure may explain the lower dimensionality of the random Gaussian process relative to the standard input; we also believe that it would increase the complexity of the transformation to abstract representations, which may explain the lower regression generalization performance for the random Gaussian process inputs as well.

Finally, we ask whether the multi-tasking model can produce abstract representations from a third kind of input, one chosen to mimic the structure of a population of highly local Gaussian receptive fields (RF), which are thought to be used to encode many different kinds of stimuli across different sensory systems[18–22]. Here, we construct a representation of a *D* = 2 latent variables using Gaussian receptive fields (fig. S4a, left) which induces a highly curved geometry in population space (fig. S4a, right). Then, we test whether or not the multi-tasking model can recover abstract representations from this highly nonlinear format. While this format is lower dimensional than the standard input used previously, it is constructed to have no global structure (i.e., each neuron responds only to a local region of latent variable space). Due to this lack of global structure, while almost any binary classification of the input space can be implemented with high accuracy, the classifier generalization performance is near chance (fig. S4b, left). The regression metric follows this same pattern: While the standard performance of a linear regression is relatively high, the regression generalization performance is at chance (fig. S4b, right). Training the multi-tasking model on these inputs produces partially abstract representations: The classifier generalization performance becomes high while the regression generalization performance remains at chance even for many tasks (fig. S4c).

### 2.7 The multi-tasking model produces abstract representations from image inputs

Given the previous results showing that the multi-tasking model produces only partially abstract representations from highly tangled or highly local inputs (i.e., the low length scale random Gaussian process or RF inputs explored in the previous section), we next asked whether the multi-tasking model would produce fully (i.e., high classification and regression generalization performance) or partially (i.e., only high classifier generalization performance) abstract representations of the image inputs often used to study disentangling in the machine learning literature (e.g., [32]): A chair image dataset that includes 3D rotations[33] and a simple 2D shape dataset[34] (fig. 5a). First, we pre-process the images using a deep network trained to perform object recognition (see *Preprocessing using a pre-trained network* in *Methods*). These networks have been shown to develop representations that resemble those found in brain regions like the inferotemporal cortex (ITC)[35], at the apex of the primate ventral visual stream. Then, we proceeded as before, using a two layer network to learn several distinct classification tasks (fig. 5b, and see *The image datasets* in *Methods* for more details).

**Figure 5:**
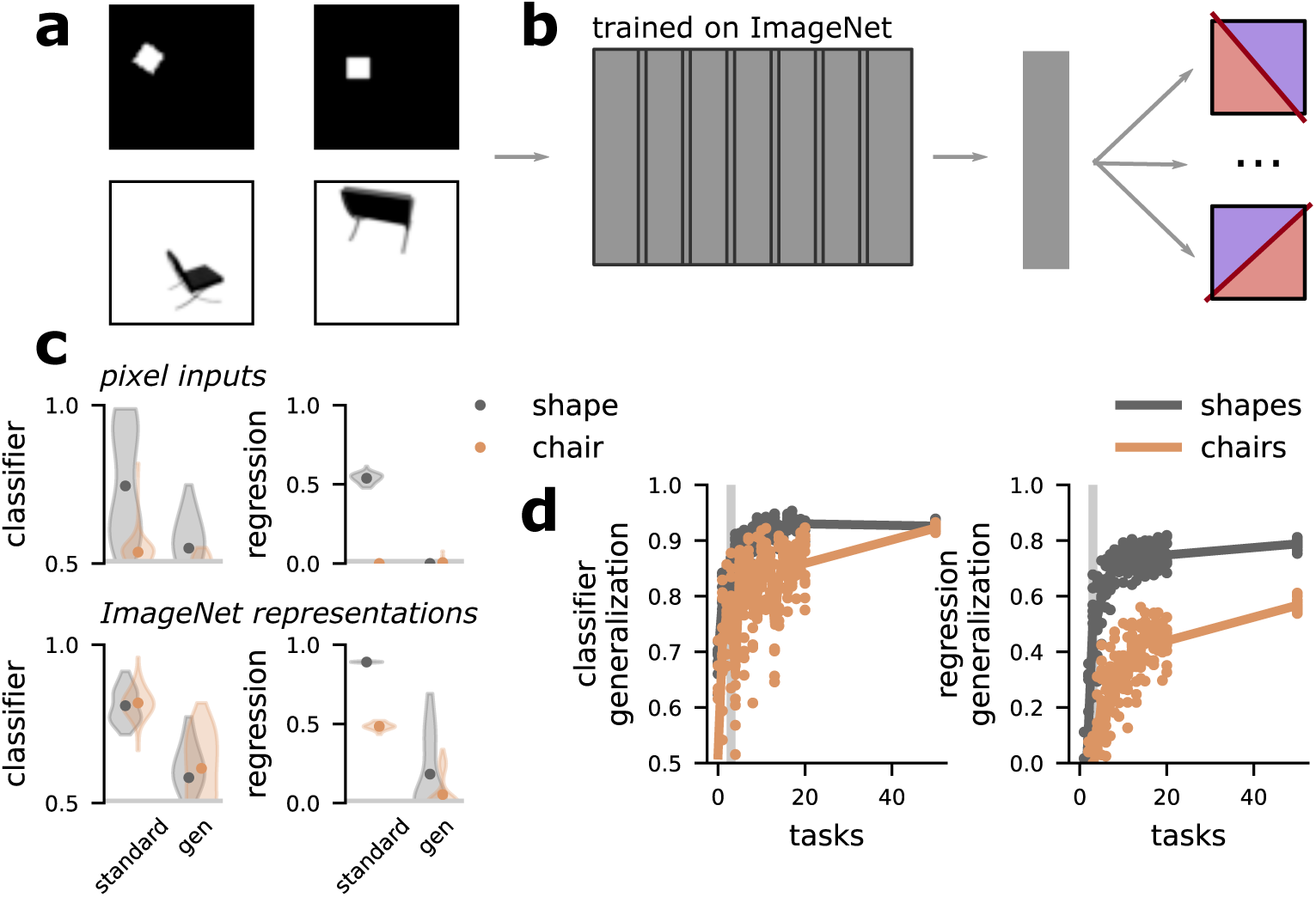
The multi-tasking model produces abstract representations from image inputs. **a** Examples from the 2D shape dataset (top) and chair image dataset (bottom). **b** Schematic of modified model. The multi-tasking model now begins with a networked pre-trained on the ImageNet challenge, followed by a few additional layers of processing before learning binary tasks as before (see *Pre-processing using a pre-trained network* in *Methods*). **c** The classifier (left) and regression (right) generalization performance when applied to the shape and chair image pixels (top) or ImageNet representations (bottom). **d** The classifier (left) and regression (right) generalization performance of the multi-tasking model trained as described in **a** on the shape and chair images.

The images in both datasets are described by three continuous parameters and one categorical variable. The chair images have continuous horizontal position, vertical position, and azimuthal rotation variables, along with the categorical chair type variable. The 2D shape images are described by continuous horizontal position, vertical position, and scale variables, along with the categorical shape type variable. For both datasets, the tasks learned by the model depend only the continuous variables, not on the categorical variables.

In both datasets, the pixel-level images (fig. 5c, top) and the representations produced by the pretrained network alone (fig. 5c, bottom) are non-abstract. However, the representations produced by the multi-tasking model are abstract, and show strong classifier generalization performance and moderately high regression generalization performance (fig. 5d). Thus, the multi-tasking model can produce fully abstract representations from representations of objects similar to those observed in the brain.

### 2.8 The multi-tasking model learns abstract representations using reinforcement learning

In all of the previous cases, we have used supervised learning to train the multi-tasking model. While this is widely used in machine learning and has been shown to produce representations that resemble those found in the brain in many cases[35–37], the information used to train the network during supervised training is qualitatively different from the information that would be received by a behaving organism performing multiple tasks. Here, we confirm that the multi-tasking model still produces abstract representations when trained using reinforcement learning.

We use a modified version of the deep deterministic policy gradient (DDPG)[38] reinforcement learning framework to train our networks. In this setup, there are two networks: an actor network, which is trained to take a stimulus and produce the action (or set of actions) that will maximize reward (fig. 6a, left) – this is directly analogous to the full multi-tasking model as previously described. In reinforcement learning, the actor cannot be trained directly from the gradient between the produced and correct actions. So, instead, a second network, referred to as the critic, is created, which is trained to predict the reward outcome from an observation and a potential action (fig. 6a, right). The critic network is trained to accurately predict the reward that results from a stimulus-action pair. Then, the actor network is trained to produce actions that lead to predicted reward (see *The reinforcement learning multi-tasking model* in *Methods* for more details). Here, we create a critic network for each of the tasks that the reinforcement learning multi-tasking model is trained to perform.

**Figure 6:**
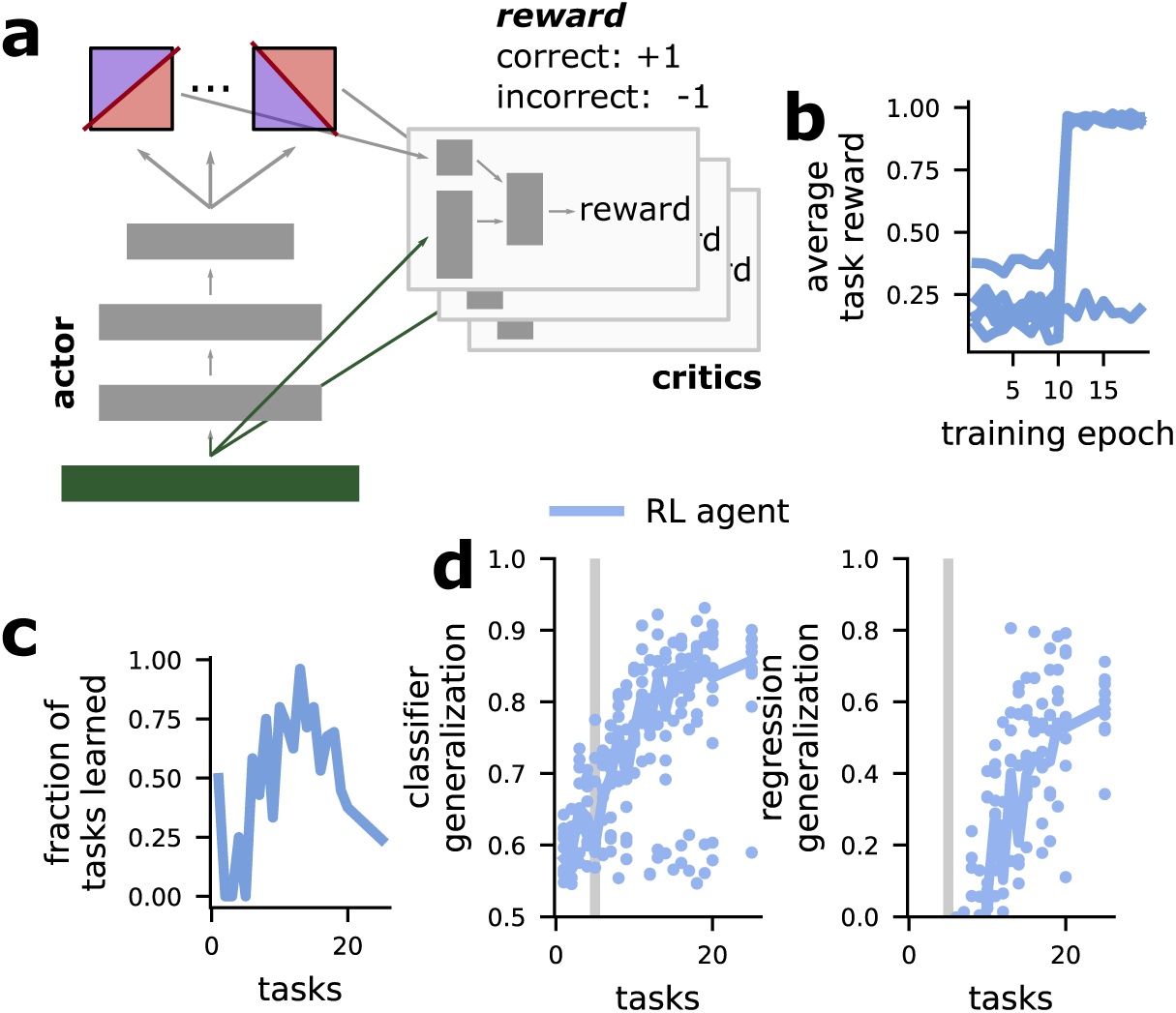
The multi-tasking model produces abstract representations when trained with reinforcement learning. **a** Schematic of the reinforcement learning multi-tasking model, using the deep deterministic policy gradient approach. **b** The performance of the network on different tasks over the course of training. Note the sharp transitions between near-chance performance (avg reward = 0) and near perfect performance (avg reward = 1). **c** The fraction of tasks learned such that avg reward *>* .8 for different numbers of trained tasks. **d** The classifier (left) and regression (right) generalization performance.

While the reinforcement learning multi-tasking model learns the tasks less reliably than the supervised multi-tasking model (fig. 6b, c), it still produces fully abstract representations for around ten trained tasks (*D* = 5, fig. 6d). Interestingly, while some tasks are not successfully learned during the allotted training time, the learned tasks transition from near-chance performance to near-perfect performance within just a few training epochs (fig. 6b). This suggests that additional hyperparameter tuning could potentially improve task learning consistency and push the representations to be even more strongly abstract with fewer trained tasks.

## 3 Discussion

We demonstrate that requiring a feedforward neural network to perform multiple tasks reliably produces abstract representations. Our results center on artificial neural networks; however, we argue that abstract representations in biological neural systems could be produced through the same mechanism, as behaving organisms often need to multi-task in the same way as we have modeled here. We show that the learning of these abstract representations is remarkably reliable. They are learned even for heterogeneous classification tasks, stimuli with partial information, in spite of being required to learn additional non-latent variable aligned tasks, and for a variety of periodic, highdimensional, and image inputs. Finally, we show that the multi-tasking model develops abstract representations even when trained with reinforcement rather than supervised learning. Overall, this work provides insight into how the abstract neural representations characterized in experimental data may emerge: Through the multiple constraints and complexity induced by naturalistic behavior.

We train our models to perform different binary classifications of latent variables as a proxy for different behaviors. This is, of course, a highly simplified approach. While feedforward binary classification most closely matches rapid objection recognition or, for example, go or no-go decisions, it does not provide an accurate model of behaviors that unfold over longer timescales. While most the experimental work that shows abstract representations in the brain[5–7, 10] and other models that produce abstract representations in machine learning systems[12, 15, 16] have taken a static view of neural activity, network dynamics could play a role in establishing and sustaining abstract representations. While some work has shown that training recurrent neural networks to perform multiple dynamic tasks leads to shared representations of common task features, the abstractness of these representations has not been quantified[24]. Future work will probe to what degree our findings here generalize to networks trained to perform dynamic tasks.

### 3.1 Other methods for quantifying abstractness

Our method of quantifying abstractness in both artificial and biological neural networks has an important difference from some previously used methods[7]. In particular, an influential model for creating disentangled representations in machine learning, the *β* variational autoencoder (*β*VAE), attempts to isolate the representation of single latent variables to single units in the network[12]. Directly applied to neural data, this leads to the prediction that single neurons should represent single latent variables in abstract representations[7]. Our framework differs in that generalization performance depends on the geometry of the representations at the population level and it is unaffected by whether single neuron activity corresponds to a single latent variable, or to a linear mixture (i.e., a weighted sum) of all the latent variables. Given the extensive linear and non-linear mixing observed already in the brain[1, 5, 6, 39], we believe that this flexibility is an advantage of our framework for detecting and quantifying the abstractness of neural representations. Further, we believe that searching for abstract representations using techniques that are invariant to linear mixing will reveal abstract representations where they may not have been detected previously – in particular, a representation can provide perfect generalization performance without having any neurons that encode only a single latent variable, and thus such a representation would not be characterized as abstract by many machine learning abstraction or disentanglement metrics.

### 3.2 Predictions

For experimental data, our findings predict that an animal trained to perform multiple distinct tasks on the same set of inputs will develop abstract representations of the latent variable dimensions that are used in the tasks. In particular, if the tasks only rely on three dimensions from a five dimensional input, then we expect strong abstract representations of those three dimensions (as in fig. S2c, d), but not of the other two. We expect all of the dimensions to still be represented in neural activity, however – we just do not expect them to be represented abstractly. Once this abstract representation is established through training on multiple tasks, if a new task is introduced that is aligned with these learned latent variables, we expect the animal to be able to learn and generalize that task more easily than a task that relies on either the other latent variables or is totally unaligned with the latent variables (as the grid tasks above). That is, we expect animals to be able to take advantage of the generalization properties provided by abstract representations that we have focused on throughout this work, as suggested by previous experimental work in humans[28].

A recent study in which human participants learned to perform two tasks while in a functional magnetic resonance (fMRI) scanner provides some evidence for our predictions[9]. The representations of a high-dimensional stimulus with two task-relevant dimensions (one which was relevant in each of two contexts) were studied in both the fMRI imaging data and in neural networks that were trained to perform the two tasks (the setup in this work is similar to certain manipulations in our study, particularly to the partial information case shown in fig. S2a, b). They find that the representations developed by a neural network which develops rich representations (similar to abstract representations in our parlance) are more similar to the representations in the fMRI data than neural networks that develop high-dimensional, non-abstract representations. This provides evidence for our central prediction: That abstract representations emerge through multiple task learning. However, the conditions explored in the human and neural network experiments in the study were more limited than those explored here. In particular, only two tasks were performed, the stimulus encoding was less nonlinear than in our studies, and the tasks were always chosen to be orthogonal. Thus, further work will be necessary to determine the limits of our finding in real brains.

Several additional predictions can be made from our results with the grid tasks, which showed that learning many random, relatively uncorrelated tasks both does not lead to the development of abstract representations alone, but also does not interfere with abstract representations that are learned from a subset of tasks that are aligned with the latent variables. First, if an animal is trained to perform a task analogous to the grid task, then we do not expect it to show abstract representations of the underlying latent variables – this would indicate that latent variables are not inferred when they do not support a specific behavior. Second, we predict that an animal trained to perform some tasks that are aligned to the latent variables as well as several (potentially more) non-aligned grid task analogues will still develop abstract representations. Both of these predictions can be tested directly through neurophysiological experiments as well as indirectly through behavioral experiments in humans (due to the putative behavioral consequences of abstract representations[28]).

### 3.3 Conclusions

Overall, our work indicates that abstract representations in the brain – which are thought to be important for generalizing knowledge across contexts – emerge naturally from learning to perform multiple categorizations of the same stimuli. This insight helps to explain previous observations of abstract representations in tasks designed with multiple contexts (such as [5]), as well as makes predictions of conditions in which abstract representations should appear more generally.

## Acknowledgments

We thank Mattia Rigotti, Nicolas Masse, and Matthew Rosen for comments on an earlier version of this manuscript. This work was supported by the following grants: Simons Foundation 542983SPI, Neuronex NSF 1707398, and Gatsby Charitable Foundation GAT3708. We also acknowledge computing resources from Columbia University’s Shared Research Computing Facility project, which is supported by NIH Research Facility Improvement Grant 1G20RR030893-01, and associated funds from the New York State Empire State Development, Division of Science Technology and Innovation (NYSTAR) Contract C090171, both awarded April 15, 2010.

## Author contributions

WJJ and SF conceived of the project and developed the simulations. WJJ performed the simulations and analytical calculation. WJJ analyzed the simulation results and made the figures. WJJ and SF wrote and edited the paper.

## M1 Methods

### M1 Code

All of our code for this project is written in Python, making extensive use of TensorFlow[1] and the broader python scientific computing environment (including numpy[2], scipy, matplotlib, and scikit-learn[3]).

The code is available at the following link: https://github.com/wj2/disentangled

### M2 Abstraction metrics

Both of our abstraction methods quantify how well a representation that is learned in one part of the latent variable space (e.g., a particular context) generalizes to another part of the latent variable space (e.g., a different context). To make this concrete, in both metrics, we train a decoding model on representations from only one – randomly chosen – half of the latent variable space and test that decoding model on representations from the non-overlapping half of the latent variable space.

#### M2.1 The classifier generalization metric

First, we select a random balanced division of the latent variable space. One of these halves is used for training, the other is used for testing. Then, we select a second random balanced division of the latent variable space that is orthogonal to the first division. One of these halves is labeled category 1 and the other is labeled category 2. As described above, we train a linear classifier on this categorization using 500 training stimuli from the training half of the space, and test the classifier’s performance on 500 stimuli from the testing half of the space. Thus, chance is set to .5 and perfect generalization performance is 1.

#### M2.2 The regression generalization metric

As above, except we train a linear ridge regression model to read out all *D* latent variables using 500 sample stimulus representations from the training half of the space. We then test the regression model on 500 stimulus representations sampled from the testing half of the space. We quantify the performance of the linear regression with its *r*^2^ value:

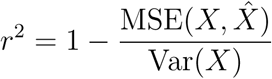

where *X* is the true value of the latent variables and 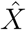 is the prediction from the linear regression. Because the MSE is unbounded, the *r*^2^ value can be arbitrarily negative. However, chance performance is *r*^2^ = 0, which would be the performance if the linear regression always predicted the mean of *X*, and *r*^2^ = 1 indicates a perfect match between the true and predicted value.

### M3 Non-abstract input generation

In the main text, we use two methods for generating non-abstract inputs from a *D*-dimensional latent variables. We have also performed our analysis using several other methods, which we also describe here.

#### M3.1 Participation ratio-maximized representations

We train a symmetric autoencoder (layers: 100, 200 units) to maximize the participation ratio[4] in its 500 unit representation layer. The participation ratio is a measure of embedding dimensionality that is roughly equivalent to the number of principal components that it would take to capture 80 % of the total variance. The autoencoder ensures that information cannot be completely lost, while the participation ratio regularization ensures that the representation will have high-embedding dimension and, therefore, be non-abstract. The performance of our generalization metrics on this input representation is shown in fig. 1f.

#### M3.2 Random Gaussian process inputs

To generate the random Gaussian process inputs, we proceed through each input dimension separately. For each dimension, we we sample a single *D*-dimensional function from the prior of a Gaussian process with a radial basis function kernel of length scale *l*. Then, the full input is simply the vector of all of these input dimensions.

We use the implementation of Gaussian processes provided in scikit-learn[3]. In particular, we initialize a Gaussian process with the above kernel, then take a sample scalar output from the Gaussian process prior distribution for a selection of 500 random points. Then, we freeze this function in place by fitting the Gaussian process to reproduce this output sample from the same set of input points.

#### M3.3 Receptive field-style representations

This input transformation is constructed to be analogous to the receptive field representations observed in many early sensory areas[5–9]. In particular, where single units respond most strongly for a particular conjunction of the *D* latent variables, and their response falls off exponentially as distance from that center point increases.

In this case, we arrange receptive field centers to tile the probability distribution of the *D*-dimensional latent variables (i.e., for Gaussian latent variables there will be more receptive field centers clustered in the center, where the probability density is higher). The width of the receptive fields is inversely related to probability density (i.e., receptive fields are wider away from the center of the latent variable distribution). The receptive fields have a Gaussian shape. That is, for a receptive field *i* with center *μ*_*i*_ and width *w*_*i*_ where both of these parameters are vectors in the *D*-dimensional latent variable space. The response of a receptive field unit *i* is given by

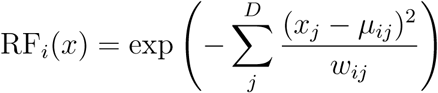

Receptive fields are particularly non-abstract, as shown by the performance of our generalization metrics directly on the receptive field representations, which is shown in fig. S4b.

### M4 The multi-tasking model

We primarily study the ability of the multi-tasking model to produce abstract representations according to our classification and regression generalization metrics. The multi-tasking model is a feedforward neural network. For figs. 2 and 3 it has the following parameters:

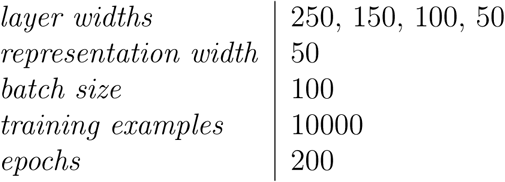

For fig. S4, everything is kept the same except the number of layers is increased:

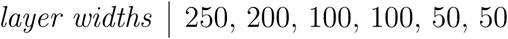

The increased number of layers improves performance in the RF case. However, for the standard input (and the images, as described below), the results are similar with only three layers (see *A sensitivity analysis of the multi-tasking model and βVAE* in *Supplement*).

#### M4.1 Full task partitions

In all cases, the models are trained to perform multiple tasks – specifically, binary classification tasks – on the latent variables. In the simplest case (i.e., fig. 2e), the task vector can be written as,

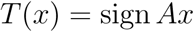

where *A* is a *P × D* matrix with randomly chosen elements.

#### M4.2 Unbalanced task partitions

For unbalanced partitions, the task vector has the following simple modification,

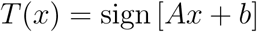

where *b* is a *P*-length vector and *b*_*i*_ *∼* 𝒩 (0, *σ*_offset_). Notice that this decreases the average mutual information provided by each element of *T* (*x*) about *x*.

#### M4.3 Contextual task partitions

We chose this manipulation to match the contextual nature of natural behavior. As motivation, we only get information about how something tastes for the subset of stimuli that we can eat. Here, we formalize this kind of distinction by choosing *P* classification tasks that each only provide information during training in half of the latent variable space.

We can write each element *i* of the contextual task vector as follows,

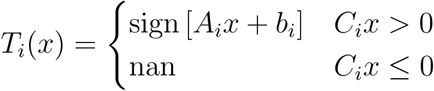

where nan values are ignored during training and *C* is a *P × D* random matrix. Thus, each of the classification tasks influences training only within half of the latent variable space. This further reduces the average information provided about *x* by each individual partition.

#### M4.4 Partial information task partitions

For contextual task partitions, the contextual information acts on particular tasks. For our partial information manipulation, we take a similar structure, but it instead acts on specific training examples. The intuitive motivation for this manipulation is to mirror another form of contextual behavior: At a given moment (i.e., sampled training example) an animal is only performing a subset of all possible tasks *P*. Thus, for a training example from that moment, only a subset of tasks should provide information for training.

Mathematically, we can write this partial information structure as follows. For each training example *x*, the task vector is given by,

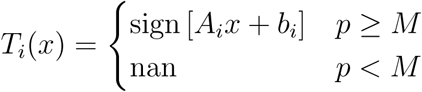

where *p* is a uniformly distributed random variable on [0, 1], which is sampled uniquely for each training example *x* and *M* is a parameter also on [0, 1] that sets the fraction of missing information. That is, *M* = .9 means that, for each training example, 90 % of tasks will not provide information.

While results are qualitatively similar for many values of *M*, in the main text we use a stricter version of this formalization: For each training sample, one task is randomly selected to provide information and the targets for all other tasks are set to nan.

#### M4.5 Grid classification tasks

The grid tasks explicitly break the latent variable structure. Each dimension is broken into *n* parts with roughly equal probability of occurring (see schematic in fig. 3a). Thus, there are *n*^*D*^ unique grid compartments, each of which is a *D*-dimensional volume in latent variable space, and each compartment has roughly equal probability of being sampled. Then, to define classification tasks on this space, we randomly assign each compartment to one of the two categories – there is no enforced spatial dependence.

#### M4.6 Random Gaussian process tasks

To generate a random Gaussian process task indexed by *i*, we sample a single *D*-dimensional function from the prior of a Gaussian process with a radial basis function kernel of length scale *l*, which we denote as 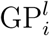. Then, to determine the category of a particular sample *x*, we evaluate the function on that category,

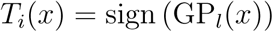

We use the implementation of Gaussian processes provided in scikit-learn[3]. In particular, we initialize a Gaussian process with the above kernel, then take a sample scalar output from the Gaussian process prior distribution for a selection of 500 random points. Then, we freeze this function in place by fitting the Gaussian process to reproduce this output sample from the same set of input points.

#### M4.7 The dimensionality of representations in the multi-tasking model

First, we consider a deep network trained to perform *P* balanced classification tasks on a set of *D* latent variables *X ∼ 𝒩* (0, *I*_*D*_). We focus on the activity in the layer just prior to readout, which we refer to as the representation layer and denote as *r*(*x*) for a particular *x ∈ X*. This representation layer is connected to the *P* output units by a linear transform *W*. In our full multi-tasking model, we then apply a sigmoid nonlinearity to the output layer. To simplify our calculation, we leave that out here. The network is trained to minimize error, according to a loss function which can be written as:

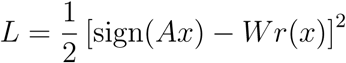

where *A* is a *P × D* matrix of randomly selected partitions (and it is assumed to be full rank). To understand how *r*(*x*) will change during training, we write the update rule for *r* (to be achieved indirectly by changing preceding weights),

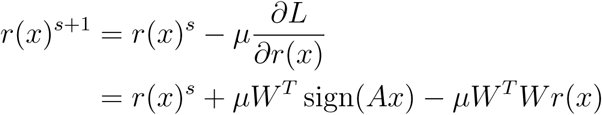

Thus, we can see that, over training, *r*(*x*) will be made to look more like a linear transform of the target function, sign(*Ax*). Next, to link this to abstract representations, we first observe that *Ax* produces an abstract representation of the latent variables. Then, we show that sign(*Ax*) has approximately the same dimensionality as *Ax*. In particular, the covariance matrix *M* = *E*_*X*_ sign(*Axx*^*T*^ *A*^*T*^) has the elements,

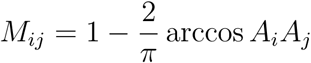

where *A*_*i*_ is the *i*th row of *A*. To find the dimensionality of sign(*Ax*) we need to find the dimensionality of *M*. First, the distribution of dot products between random vectors is centered on 0 and the variance scales as 1*/D*, where *D* is the dimensionality of the latent variables as usual. Thus, we can Taylor expand the elements of the covariance matrix around *A*_*i*_*A*_*j*_ = 0, which yields

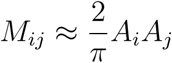

We identify this as a scalar multiplication of the covariance matrix for the linear, abstract target *E*_*X*_ [*Axx*^*T*^ *A*^*T*^]. Further, we know that the rank of this matrix is min(*P, D*). So, this implies that the matrix *M* also has rank approximately min(*P, D*). Deviations from this approximation will produce additional non-zero eigenvalues, however they are expected to be small.

Importantly, while a high dimensional *r*(*x*) can solve *P* classification tasks in a non-abstract way (for example, notice that the classification accuracy of the standard inputand RF inputs below have very high classification accuracy for random tasks yet much lower generalization performance, fig. 1f, left and fig. S4b, left), an *r*(*x*) with dimensionality min(*P, D*) will be constrained to solve the tasks in an at least partially abstract way (see *Four possibilities for representations in the multi-tasking model* in *Supplement*).

### M5 Pre-processing using a pre-trained network

When applying the multi-tasking model to image inputs, we used a deep neural network trained on the ImageNet classification task to pre-process them into a feature vector. Then, we used this representation as input to the multi-tasking model. The pre-trained network is not fine tuned, or trained, during the training of the multi-tasking model.

The parameters of the model used here are the same as the multi-tasking model except:

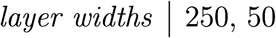

and only 300 of the unique chair types were used in the training dataset. The network we used is available here.

### M6 The image datasets

We used two standard image datasets from the machine learning literature. In both cases, we considered a subset of the total number of features, explained below.

#### M6.1 The 2D shapes dataset

This dataset consists of white 2D shapes on a black background[10]. The features are horizontal and vertical position, 2D rotation, scale, and shape type. We do not train tasks on either the rotation or shape type variables. We exclude rotation due to its periodicity and shape type because it is a categorical variable. All values of both variables are still included in the training dataset, they are simply not used in the classification tasks.

#### M6.2 The 3D chair dataset

This dataset consists of images of different styles of chairs on a white background[11]. The original features are azimuthal rotation, pitch, and chair style. We augment these features to include horizontal and vertical position by translating the image using coordinates sampled from a normal distribution and truncated when the chair portion of the image begins to wrap around the edges. We exclude pitch, a subset of azimuthal rotations, and chair styles from task training. We exclude pitch because it has only two values in the dataset and chair style because it is a categorical variable. Both are still included in the training data. We exclude a subset of azimuthal rotations to break the periodicity of the variable, which allows us to treat azimuthal rotation as a continuous, non-periodic variable.

### M7 The reinforcement learning multi-tasking model

We adapt the deep determinisitc policy gradient (DDPG)[12] to train the multi-tasking model. In particular, we train a single actor network, which is tasked with taking in a stimulus and producing an action. The action is the categorization of that stimulus on each of the *P* trained tasks. To provide supervision to this actor network, we train independent critic networks for each task, which take in the stimulus and the action produced by the actor and attempt to predict the reward that will be received from that pair. These critic networks are trained with respect to the actual reward received, and the predicted reward from the critic is used to train the actor network. Otherwise, we follow the standard DDPG approach as described in [12].

The correct action for a categorization task was *−*1 for one category and 1 for the other category. An action was rewarded if the network produced positive activity greater than the reward / punishment threshold for the former and negative activity greater than that threshold for the latter. If the activity was greater than that threshold but with the wrong sign, then the network received a punishment (i.e., negative reward). Otherwise, no reward or punishment was received.

The parameters of the reinforcement learning multi-tasking model are:

|  |  |
| --- | --- |
| *actor layer widths* | 250, 150, 50 |
| *representation width* | 50 |
| *critic stimulus layer widths* | 10, 5 |
| *critic action layer widths* | 1 |
| *critic shared layer widths* | 5 |
| *batch size* | 200 |
| *training epochs* | 19953 |
| *initial batches* | 20000 |
| *reward / punishment threshold* | $\pm.33$ |

### M8 The *β*VAE

The *β*VAE is an autoencoder designed to produce abstract (or, as referred to in the machine learning literature, disentangled) representations of the latent variables underlying a particular dataset[13]. The *β*VAE is totally unsupervised, while the multi-tasking model receives the supervisory task signals. Abstract representations are encouraged through tuning of the hyperparameter *β*, which controls the strength of regularization in the representation layer, which penalizes the distribution of representation layer activity for being different from the standard normal distribution. In fig. S3, the *β*VAE is trained with the same parameters as given in section M4 – the layers are replicated in reverse for the backwards pass through the autoencoder. For fig. S4, the parameters are as described in section M4. In both cases, instead of fitting models across different numbers of partitions, we fit the models with different values chosen for *β*.

For fig. S5, parameters for the *β*VAE are as described in section M9. We also explored numerous other architectures for the *β*VAE in that figure, but never obtained qualitatively or quantitatively better results.

### M9 The generative multi-tasking model

To move our multi-tasking model into a generative context, we simply add a series of layers connected to the representation layer that are trained to reproduce the original stimulus. Our objective function then has two parts: The first is to satisfy the training classification tasks and the second is to reconstruct the original input, as with a traditional autoencoder. The generative multi-tasking model was trained on the 2D shapes dataset (which were resized to be 32 *×* 32 images) with the following parameters,

|  |  |
| --- | --- |
| *layer widths* | 128x2x2, 128x2x2, 512, 256, 128, 128 |
| *representation width* | 50 |
| *batch size* | 30 |
| *training examples* | 100000 |
| *epochs* | 200 |

For the reconstruction part of the model, the given layer list is reversed.

## S1 Supplement

### S1 Four possibilities for representations in the multi-tasking model

We consider four distinct kinds of representations that could support the simultaneous performance of *P* classification tasks as formalized in the multi-tasking model. First, the high dimensional standard input could be preserved, or only weakly tuned – in particular, recall that high classification performance is already achieved on the standard input for random tasks (fig. 1f, left “standard”) even though it is not abstract (fig. 1f, left “gen”). Second, the representation could split along *P* separate dimensions of population activity, where each dimension corresponds to one of the *P* distinct tasks (fig. S1b, left). Third, the representation could consist only of an approximately *D*-dimensional sphere (or circle, in two dimensions), which exploits the correlation structure in the *P* different tasks (that is, when *P > D*, the outcomes from some pairs of tasks are necessarily correlated with each other; fig. S1b, middle). This second type of representation would have high classifier generalization performance but low regression generalization performance: That is, it is partially abstract in that it would recover the angular structure of the latent variables (as necessary for the *P* classification tasks), but not their magnitude (as this information is not necessary to solve the *P* tasks). Fourth, a fully abstract representation of the latent variables could be recovered. That is, the representation could recover both the angular structure of the latent variables, as in the second possibility, and their magnitude (fig. S1b, right). This would occur only if the multitasking model does not discard information about the stimuli that is not necessary for satisfying the tasks, but which is also not explicitly trained to discard. Surprisingly, as we will see, this fourth form of representation is most common in our trained networks, even for more disordered tasks than we have described so far.

### S2 Comparing the multi-tasking model with the unsupervised *β*VAE

We compare the level of abstraction of the representations learned by the multi-tasking model to those learned by an auto-encoder that is designed to produce abstract representations. In particular, the *β*-variational autoencoder (*β*VAE) is the current state-of-the-art for unsupervised disentangling of latent variables[1] (and it has many variations[2, 3]). It is designed around a hyperparameter, *β*, that is thought to control the trade-off between the abstractness of the representations in the latent variables and reconstruction error for output from the auto-encoder. That is, increasing *β* is understood to increase the level of abstraction in the *β*VAE representation layer, while decreasing the quality of of reconstruction of the original input representation.

Using the same architecture as in our multi-tasking model, we trained *β*VAEs to disentangle the same set of latent variables as in our other experiments. Applying the same two metrics as to our other models, we found that the *β*VAE produces moderately abstract representations, as quantified by the classifier generalization metric – though classifier generalization performance does not saturate to the same level as for the multi-tasking model. The *β*VAE does not produce high regression generalization performance for any choice of *β* that we tested. Because the multi-tasking model receives binary supervisory input and the *β*VAE does not receive any supervisory input at all, it is not particularly surprising that the multi-tasking model develops more abstract representations. However, we believe the contrast is still informative, as it indicates that abstract representations are unlikely to emerge by chance or without explicit training on tasks that are at least coarsely related to the latent variables of interest (and see [4]). Further, this multi-tasking approach to producing abstract representations is less sensitive to changes in model and input parameters than the *β*VAE (see *A sensitivity analysis of the multi-tasking model and βVAE* in *Supplement*). This further indicates the feasibility of the multi-tasking approach in conditions similar to those found in the brain.

**Figure S1:**
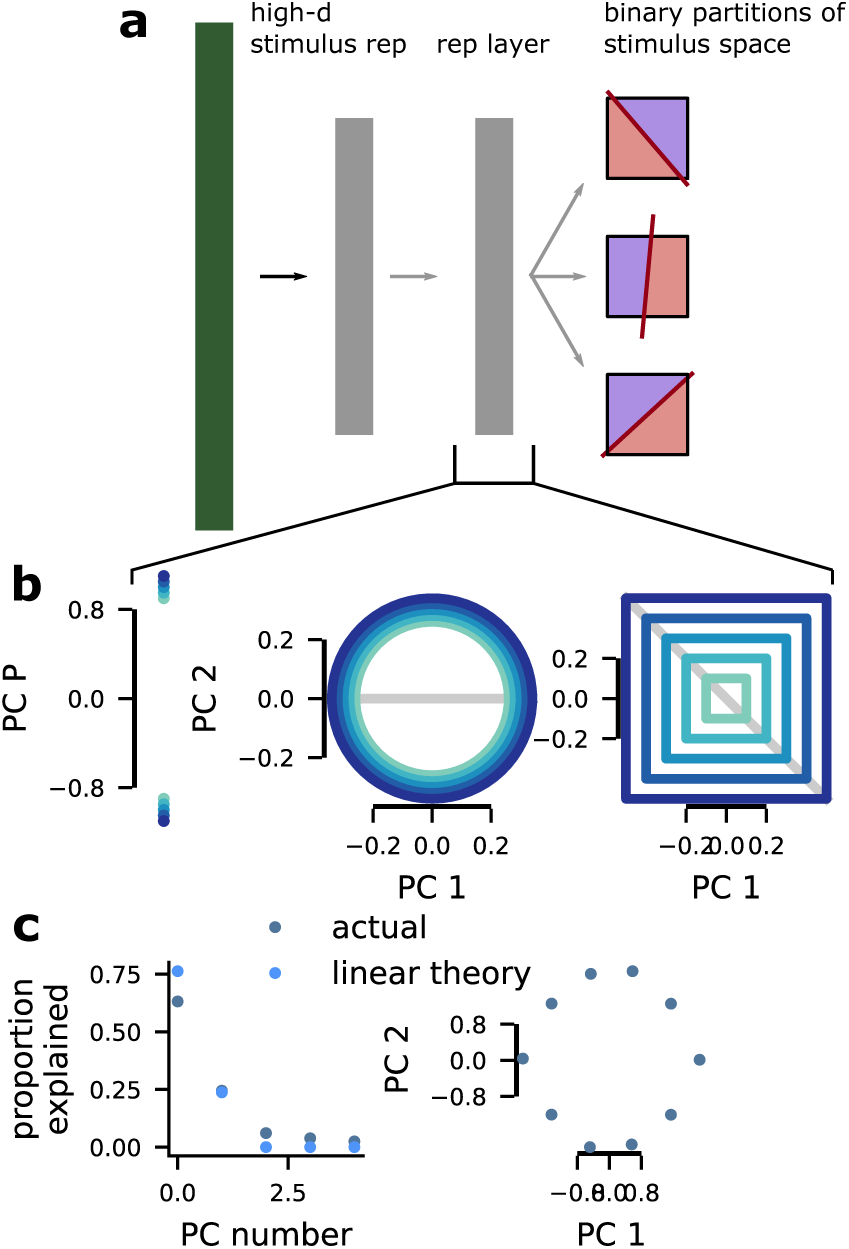
Possibilities for learned representations. **a** Schematic of the multi-tasking model. It receives an entangled stimulus representation (as shown in fig. 1e, left) and learns to perform *P* binary classifications of the latent variables. We study the representations that this induces in the layer prior to the output: the representation layer. **b** Different possible solutions the network could learn. (left) The network could learn a dimension for each classification task and develop binary representations along each of those *P* dimensions. (middle) The network could learn a surface that matches the dimensionality *D* of the latent variables, but discards information about magnitude; this representation would have high classifier- but low regression-generalization performance. (right) The network could learn a fully abstract, approximately *D*-dimensional representation of the latent variables. **c** The dimensionality of the representation layer will be approximately *D*-dimensional (left), as predicted by eq. (1). (right) The first two principal components of the required output of the network; this structure is consistent with both the middle and right network solutions from **b**.

### S3 Abstract structure can be learned from early sensory-like representations

While we explored highly nonlinear representations in the main text using low length scale random Gaussian process inputs, we also explored a special case: Gaussian receptive field inputs, which may be present in early sensory and other areas.

**Figure S2:**
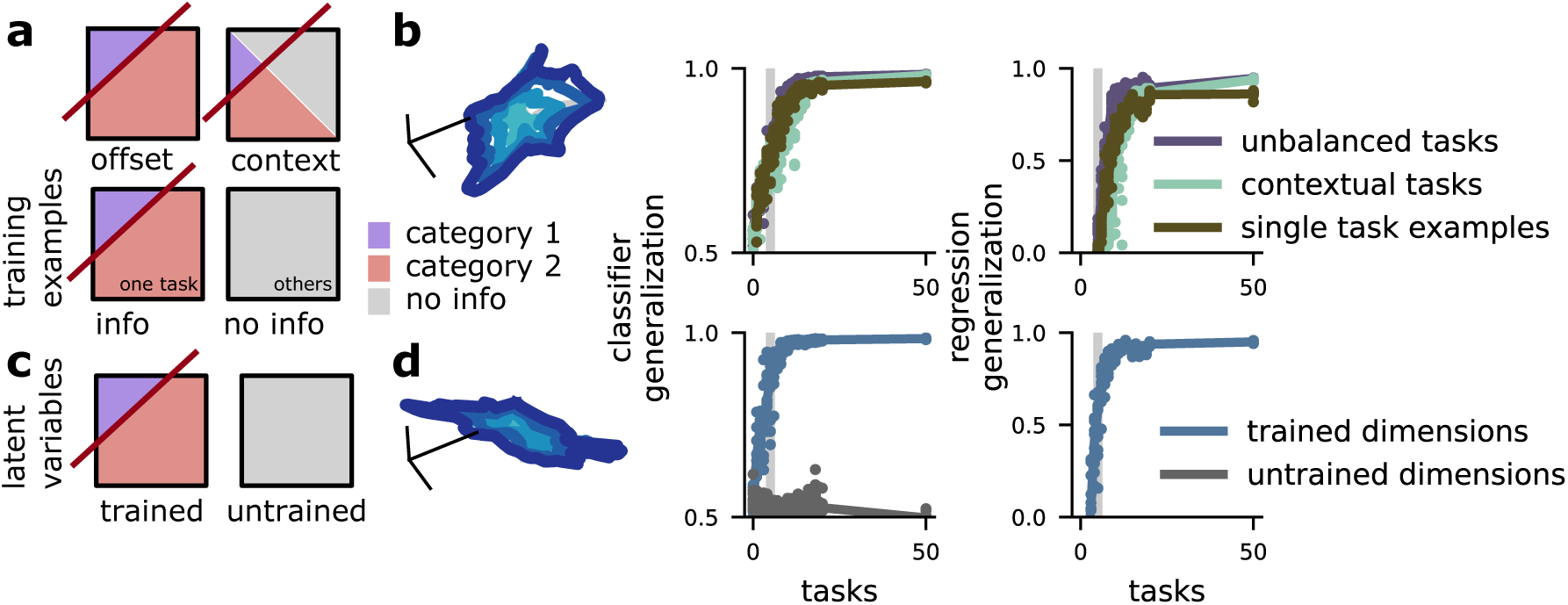
Abstract representations emerge for heterogeneous tasks, and in spite of high-dimensional grid tasks. **a** Schematics of different task manipulations. **b** (left) Visualization of the representations developed for contextual tasks *P* = 25. (middle) Classifier generalization performance. (right) Regression generalization performance. **c** Schematic showing the training scheme: A subset of latent variables are involved in tasks (left), the rest of the latent variables are not (right). **d** (left) Visualization of the trained latent variable representations. (middle) Classifier generalization performance for the trained and untrained latent variable dimensions. (right) Regression generalization performance for the trained and untrained latent variable dimensions.

**Figure S3:**
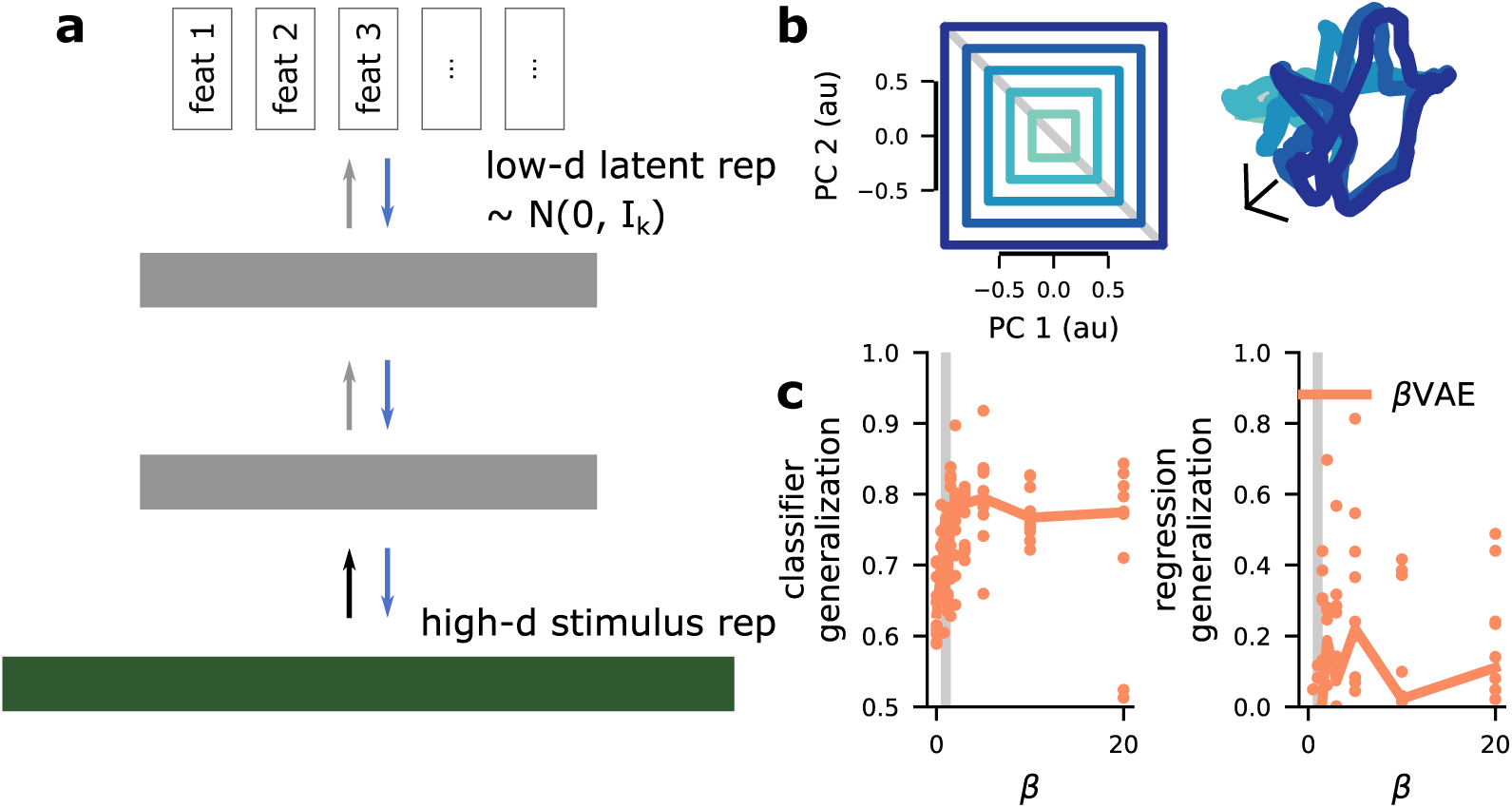
The *β*VAE does not reliably produce abstract representations. **a** A schematic of the *β*VAE. It is an autoencoder regularized to produce a low-dimensional representation in its representation layer. **b** A purely autoencoding approach with the *β*VAE is not supplied with any classification tasks (left), but does produce moderately abstract representations (right). **c** The *β*VAE produces high classifier generalization performance for a small range of *β*s (left), but does not provide high regression generalization performance for any choice of *β* that we tested (right).

Here, we construct a representation of a *D* = 2 latent variable using Gaussian receptive fields (fig. S4a, left) which induces a highly curved geometry in population space (fig. S4a, right). Then, we test whether or not the multi-tasking model can recover abstract representations from this highly nonlinear format. While this format is lower dimensional than the dimensionality-maximized input used previously, it is constructed to have no global structure (i.e., each neuron responds only to a local region of latent variable space). As a consequence of this receptive field-like format, almost any binary classification of the input space can be implemented with high accuracy, but the classifier generalization performance is near chance (fig. S4b, left). This is a consequence of the lack of global structure in the representation. The regression metric follows this same pattern: While the standard performance of a linear regression is relatively high, the regression generalization performance is at chance (fig. S4b, right).

Now, we train the multi-tasking model with these receptive field-like representations as input. We visualize the representations developed after training, as before, and show that an imperfect abstract structure is developed for a moderate number of classification tasks (fig. S4c, left). We compare this learned representation to the representation learned by the *β*VAE, given the same input and architecture (see *Comparing the multi-tasking model with the unsupervised βVAE* in *Supplement* for more details). The *β*VAE does not develop strongly abstract representations for this input – and appears to retain much of the curved structure present in the input (fig. S4c, right).

We quantify this result using our classification and regression generalization metrics. As anticipated by the visualization, the classification metric saturates performance when supplied with 8 classification tasks (fig. S4d, left, blue line). The *β*VAE did not reach above 90 % classifier generalization performance for any value of *β* that we tested (fig. S4d, left, orange line). In contrast, neither model saturates performance for the regression generalization metric (fig. S4d, right). Because the multitasking model receives binary supervisory input and the *β*VAE does not receive any supervisory input at all, it is not particularly surprising that the multi-tasking model develops more abstract representations. However, we believe the contrast is still informative, as it indicates that abstract representations are unlikely to emerge by chance or without explicit training on tasks that are at least coarsely related to the latent variables of interest (and see [4]).

**Figure S4:**
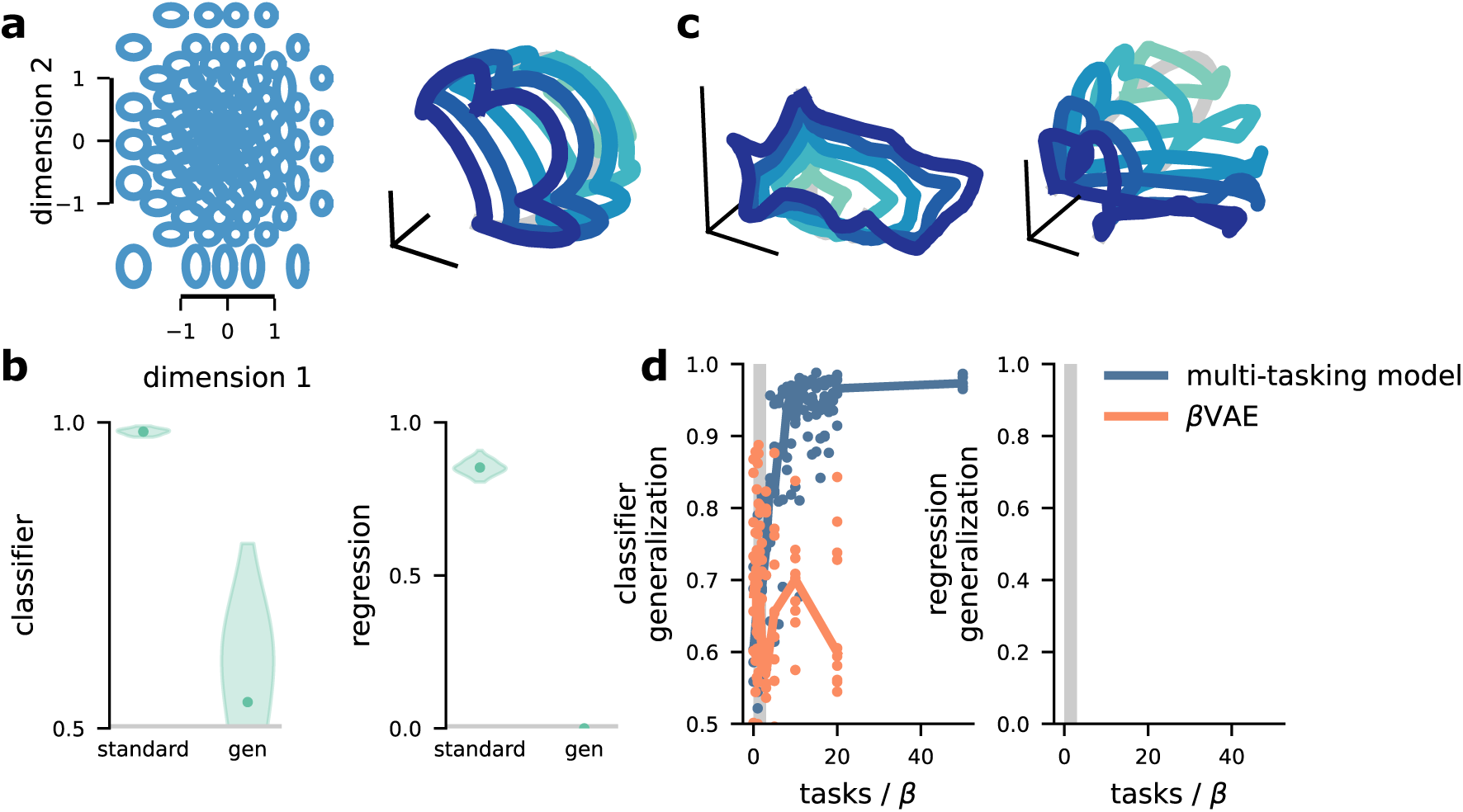
Abstract representations can be recovered even from highly nonlinear stimulus representations. **a** (left) Schematic of the receptive field inputs. They are arranged with density and RF width related to the probability density of the Gaussian inputs. (right) Low-dimensional projection of the RF representation, illustrating its high curvature. **b** Performance of the classifier-(left) and regression-metrics (right) on the RF inputs. **c** (left) Visualization of the dominant low-dimensional structure learned by the multi-tasking model. (right) Visualization of the dominant low-dimensional structure learned by the matched *β*VAE. **d** Quantification of how the abstraction of the representations learned depends on the number of classification tasksfor the multi-tasking model and *β* for the *β*VAE.

### S4 The multi-tasking model can be used as an abstract, generative model

In the main text, we show that the multi-tasking model produces abstract representations from two image datasets used in machine learning. One of the main applications of abstract (or, “disentangled”) representations in machine learning is the generation of novel images with particular latent variable values – as from the *β*VAE[5].

Here, we demonstrate that the multi-tasking model can also be used in this generative context, to produce images with expected latent variable values from the 2D shape dataset. In addition, we compare the performance of the multi-tasking model to the *β*VAE. Importantly, the multitasking model is supplied with categorical information that is related to the latent variables, as it is throughout the paper, so this comparison does not put the *β*VAE and multi-tasking model on equal footing; the *β*VAE is designed to develop abstract representations in a fully unsupervised setting. Further, we also modify the multi-tasking model, as described before, to add an autoencoder. Now, the multi-tasking model is trained to both satisfy the *P* classification tasks as well as reconstruct the original image sample to test its generative properties.

We selected one shape to be left out as a test shape, and then used the representations corresponding to the other two shapes to learn a linear regression that decodes shape scale. We then used this linear regression to generate images of one of the trained shapes at different scales (fig. S5e,f, top row). Both models retained a reasonable degree of shape structure as well as produced an increase in scale, moving from left to right in the images shown. Next, we attempted to apply the learned representation of scale to the left out shape. Again, both models produce shapes with an increase in scale (fig. S5e,f, bottom row). However, while the multi-tasking model produces images with the left out shape, the *β*VAE does not represent a differentiated shape at all. This issue with the *β*VAE has been reported before: To achieve a high level of abstraction in the representations, the *β*VAE often sacrifices precision in its reconstruction of the target image[1] (but see [2]). In contrast, the multi-tasking model produces abstract representations while still preserving its ability to reconstruct different shapes from this dataset.

**Figure S5:**
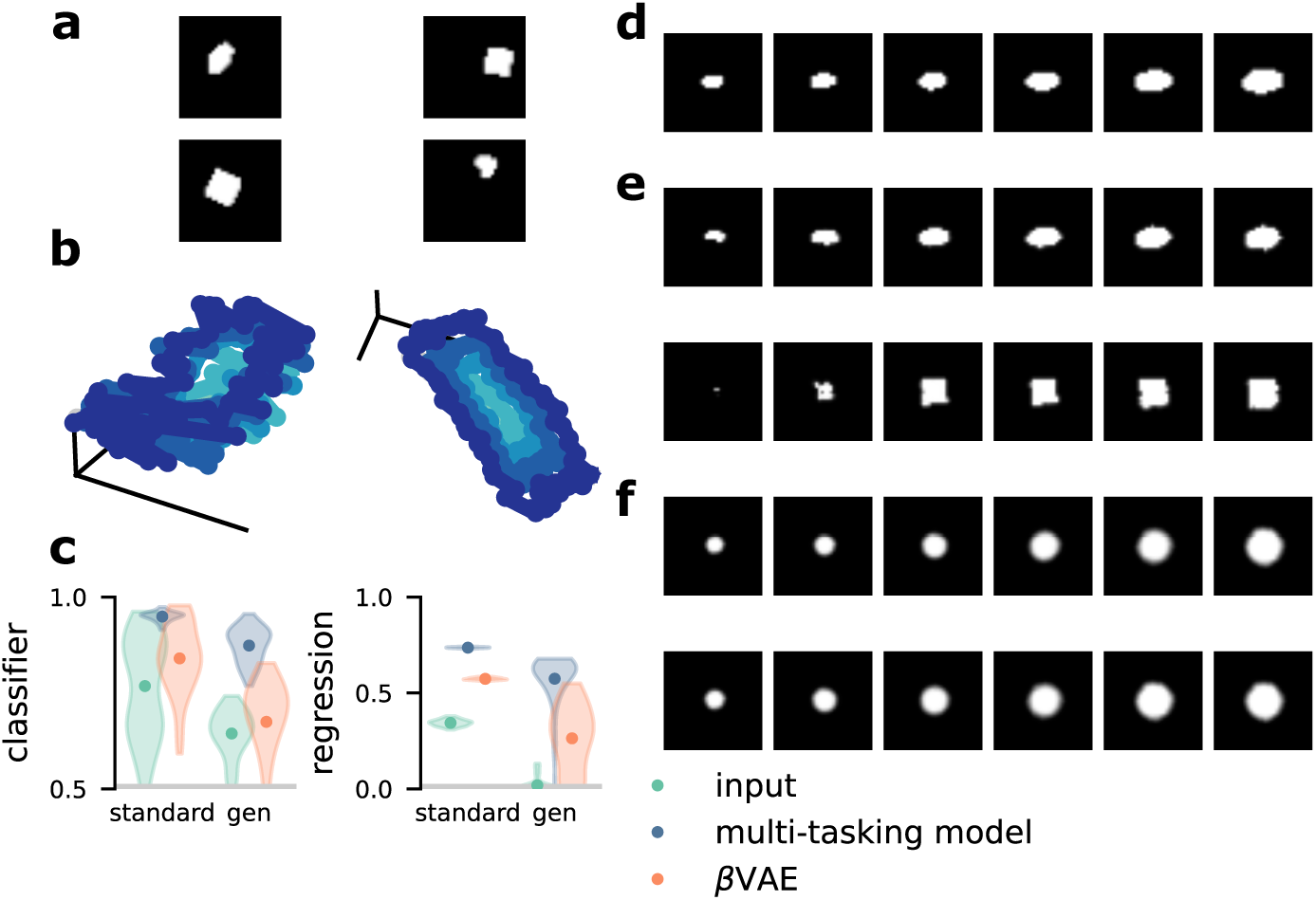
The multi-tasking model can be used for image generation. **a** Example images from the 2D shapes dataset. **b** Visualization of the image representation manifold for x- and y-position from the representation layers of the multi-tasking model (left) and the *β*VAE (right). **c** Quantification of the abstractness of the original dataset (left points), the multi-tasking model (middle points, *P* = 50), and the *β*VAE (right points, *β* = 1) according to both our classifier(left) and regression-generalization (right) metrics. In each plot, performance when training and testing on the whole stimulus set is on the left, and training and testing on separate halves is on the right; chance for both is shown by the grey line. **d** An example traversal of the scale dimension from the image set. **e** Image reconstruction for the multi-tasking model with a shape that was present in the training set (top) and that was held out from the training set (bottom). **f** The same as **e** but for the *β*VAE.

### S5 The dependence of learned abstract representations on latent variable dimensionality

For both the multi-tasking model and *β*VAE, our simulations reveal that abstract representations are more readily and consistently produced for higher-dimensional latent variables (fig. S6). We believe that this is due to a decrease in dimensionality expansion per latent dimension as *D* increases. In particular, for each of *D* = 2, 3, 4, 5, the participation ratio after expansions is approximately 200 and the participation ratio per latent variable dimension is approximately 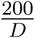. However, further work is necessary to confirm this intuition.

**Figure S6:**
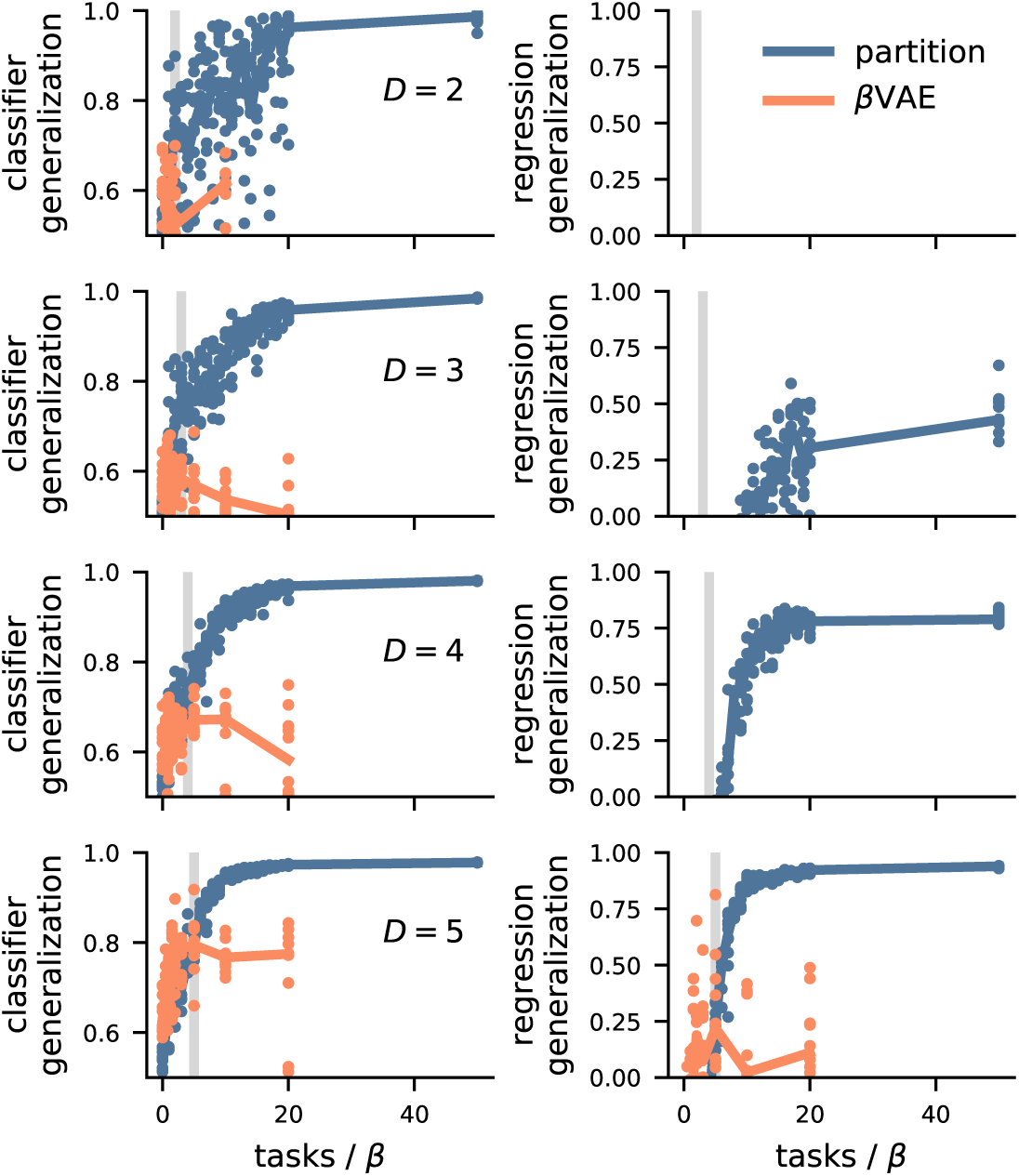
Abstraction learning depends on latent variable dimensionality. (top to bottom) Increasing latent variable dimensionality *D*, from *D* = 2 to *D* = 5 (see left inset). (left) Classifier generalization performance as a function of the number of classification tasks for the multi-tasking model and *β* for the *β*VAE. (right) Regression generalization performance as a function of the number of classification tasks for the multi-tasking model and *β* for the *β*VAE.

### S6 A sensitivity analysis of the multi-tasking model and *β*VAE

While we have focused on manipulation of the number (and kind) of classification tasks provided to the multi-tasking model and to the value of *β* for the *β*VAE, both models depend on numerous other parameter choices, which were essentially arbitrary. The parameters were held constant across the two models, but these choices can still affect the results produced by both models in different ways. To explore the dependence of our results on these other parameter choices, we performed a “multiverse” sensitivity analysis[6]. That is, for many of the parameters of our models, we chose several other similarly reasonable parameter values, and trained models with those parameters (e.g., using a tanh nonlinearity instead of the ReLU). In exploring this parameter space, we defined 7128 and 3369 distinct combinations of parameters for the multi-tasking model and *β*VAE respectively. Then, for each of these parameter combinations, we trained two models of the corresponding type and averaged their classification and regression generalization performance. The parameters varied for each were the same except for the choices of the number of classification tasks, values of *β*, and we included a version of the multi-tasking model with and without an autoencoder. To analyze these results, we fit linear models with ridge regression to account for the classification and regression generalization performance from the different parameter choices. Using only the first order version of this model (that is, without fitting interaction terms for the different parameters), the model has *r*^2^ = .62 and *r*^2^ = .70 for the multi-tasking model and *β*VAE, respectively. As expected, for the multi-tasking model, the number of classification tasks has by far the strongest effect on both classification and regression generalization performance (fig. S7a,b), though minor effects on both are produced by almost all the other parameter choices – and the regression generalization metric is strongly affected by the dimensionality of the latent variables (fig. S7b). Surprisingly, while choice of *β* does affect classification and regression generalization performance for the *β*VAE, the size of the effect is similar in size to the effects associated with many of the other parameters – and much smaller than the increase in both classification and regression generalization performance that is produced by using a tanh nonlinearity rather than a ReLU nonlinearity.

### S7 Zero-shot categorical generalization for image inputs

For the image inputs used in the main text, we also exploit the fact that they are described by a mixture of continuous and categorical features to explore other tests of generalization. In particular, while both sets of images were described by three close-to-continuous features (both: x- and y-position, chairs: 3D rotation, shapes: size), they both also had a fourth categorical feature: chair and shape type (see fig. S8a,c for examples). First, we trained the multi-tasking model using only a subset of the shapes (or chairs) and then characterized the classification and regression generalization performance using a completely novel set of shapes (chairs, fig. S8a). The shapes have high classification and regression generalization performance and the chairs have high classifier generalization performance but low regression generalization performance for this test of generalization (fig. S8b).

Then, we test another form of zero-shot generalization. As before, we train the multi-tasking model on only a subset of shapes (chairs). Then, when evaluating the classification and regression generalization performance, we train the models on that same set of shapes (chairs) in a restricted section of the latent variable space (as usual), and then evaluate the performance on those models on both the held out shapes (chairs) and the left out section of the latent variable space (fig. S8c). In this case, the shape dataset has high classification and regression generalization performance while the chair dataset has high classifier generalization performance but chance-level regression generalization performance (fig. S8d). Thus, even for this strong test of generalization to unseen images, the multi-tasking model succeeds at producing fully abstract representations of the shape dataset and partially abstract representations of the chair dataset.

---

**Figure S7:**
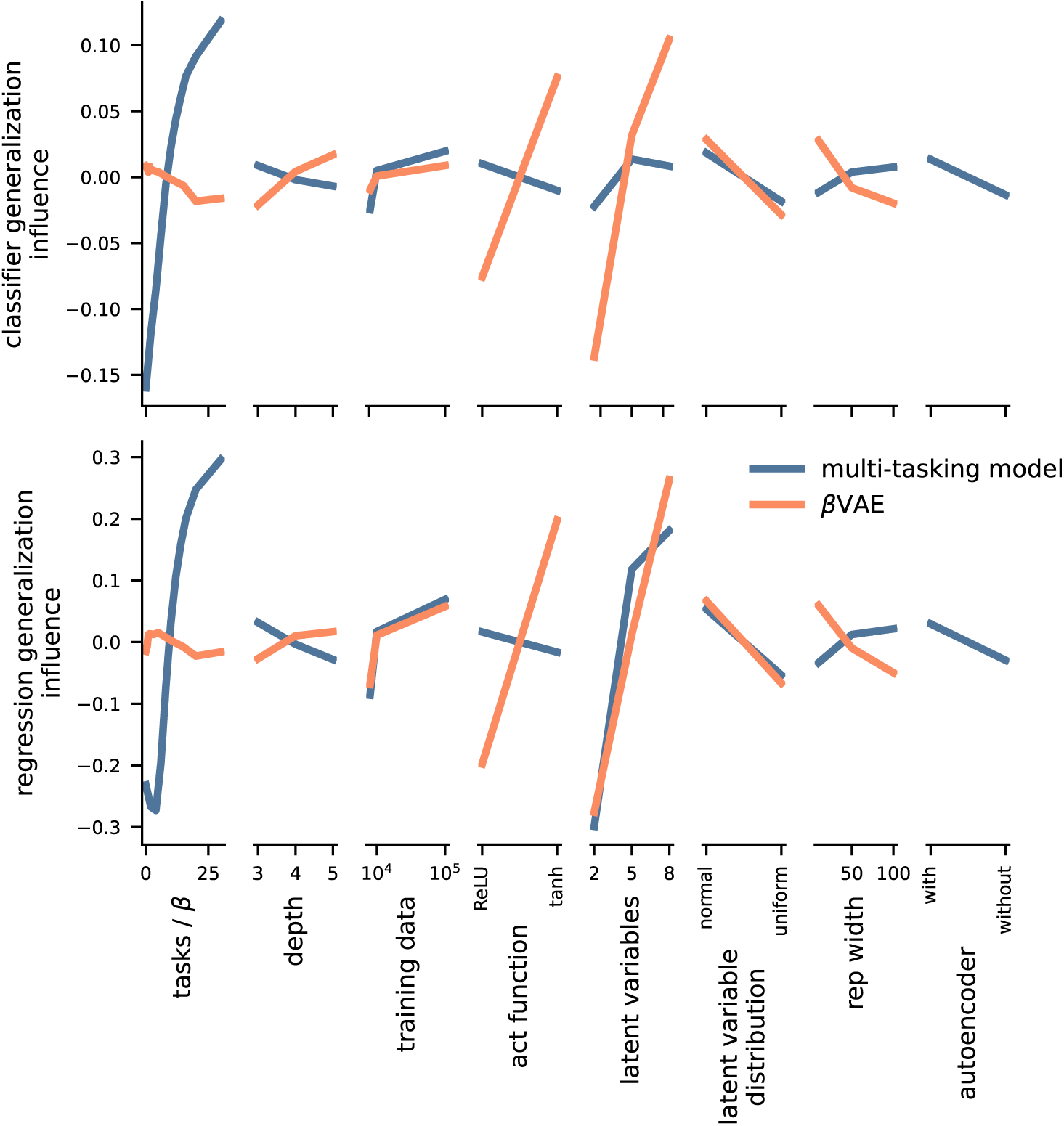
A multiverse analysis of both the multi-tasking model and *β*VAE. (top) The effects of different parameter choices on classifier generalization performance. (bottom) The effects of different parameter choices on regression generalization performance.

**Figure S8:**
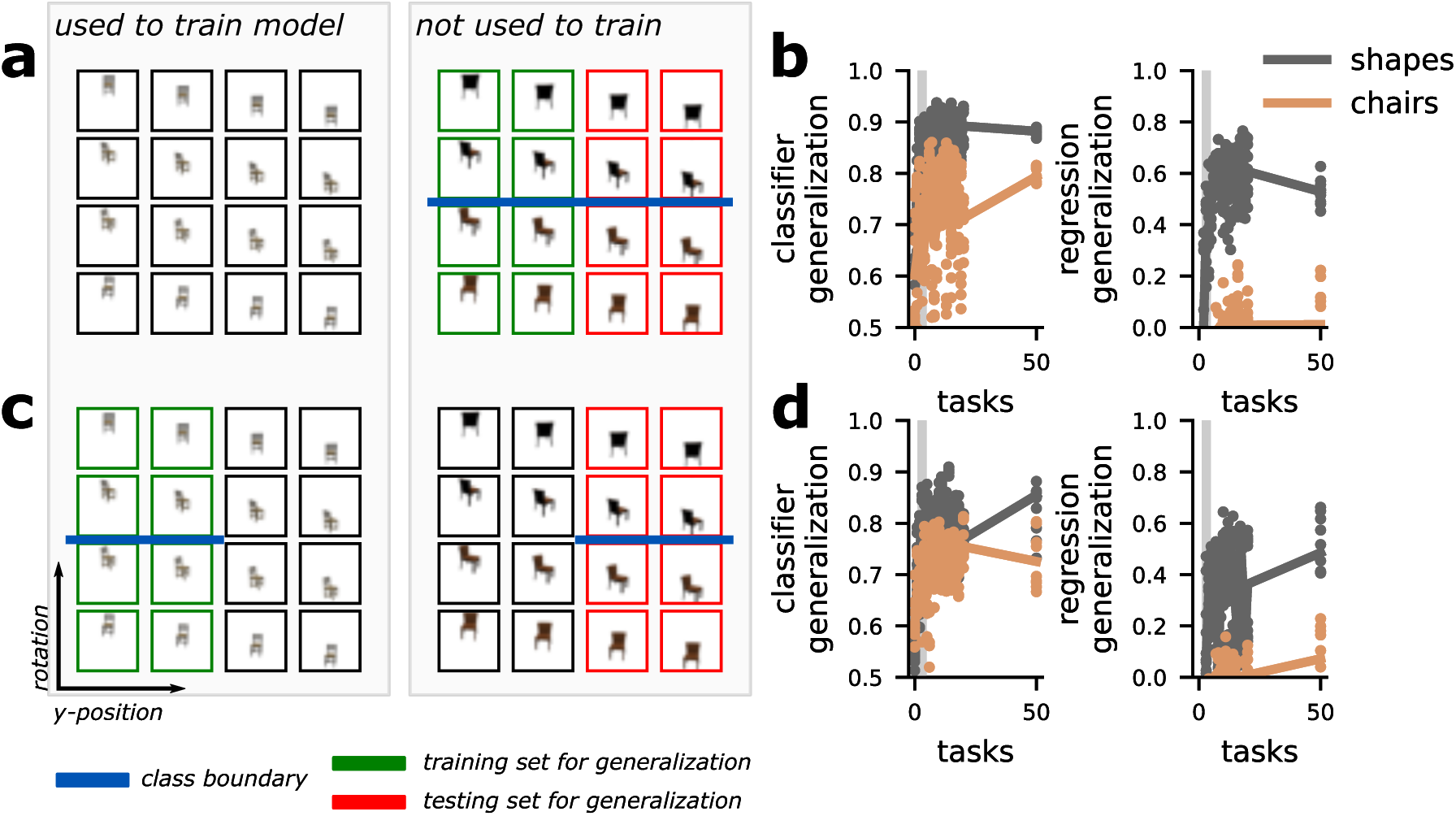
Tests of zero-shot generalization to novel images. **a** Schematic of the model training (boxes) and generalization analysis (red, green highlights) procedures for the first test of generalization. **b** Generalization performance for the training procedure shown in **a. c** Same as **a** for a second type of generalization analysis. **d** Generalization performance for the training procedure shown in **c**.

## References

1. Rigotti, M. et al. The importance of mixed selectivity in complex cognitive tasks. Nature 497, 1–6. http://dx.doi.org/10.1038/nature12160%7B%5C%%7D5Cnpapers3://publication/doi/10.1038/nature12160 (2013).

2. Fusi, S., Miller, E. K. & Rigotti, M. Why neurons mix: High dimensionality for higher cognition. Current Opinion in Neurobiology 37, 66–74. http://dx.doi.org/10.1016/j.conb.2016.01.010 (2016).

3. Stringer, C., Pachitariu, M., Steinmetz, N., Carandini, M. & Harris, K. D. High-dimensional geometry of population responses in visual cortex. Nature, 1 (2019).

4. Johnston, W. J., Palmer, S. E. & Freedman, D. J. Nonlinear mixed selectivity supports reliable neural computation. PLoS computational biology 16, e1007544 (2020).

5. Bernardi, S. et al. The geometry of abstraction in the hippocampus and prefrontal cortex. Cell 183, 954–967 (2020).

6. Chang, L. & Tsao, D. Y. The code for facial identity in the primate brain. Cell 169, 1013–1028 (2017).

7. Higgins, I. et al. Unsupervised deep learning identifies semantic disentanglement in single inferotemporal neurons. arXiv preprint 2006.14304 (2020).

8. She, L., Benna, M. K., Shi, Y., Fusi, S. & Tsao, D. Y. The neural code for face memory. bioRxiv (2021).

9. Flesch, T., Juechems, K., Dumbalska, T., Saxe, A. & Summerfield, C. Orthogonal representations for robust context-dependent task performance in brains and neural networks. Neuron (2022).

10. Sheahan, H., Luyckx, F., Nelli, S., Teupe, C. & Summerfield, C. Neural state space alignment for magnitude generalization in humans and recurrent networks. Neuron 109, 1214–1226 (2021).

11. Bengio, Y., Courville, A. & Vincent, P. Representation learning: A review and new perspectives. IEEE transactions on pattern analysis and machine intelligence 35, 1798–1828 (2013).

12. Higgins, I. et al. β-VAE: Learning basic visual concepts with a constrained variational frame-work in ICLR (2017).

13. Burgess, C. P. et al. Understanding disentangling in β-VAE. arXiv preprint 1804.03599 (2018).

14. Higgins, I., Racanière, S. & Rezende, D. Symmetry-Based Representations for Artificial and Biological General Intelligence. arXiv preprint 2203.09250 (2022).

15. Kulkarni, T. D., Whitney, W., Kohli, P. & Tenenbaum, J. B. Deep convolutional inverse graphics network. arXiv preprint 1503.03167 (2015).

16. Chen, X. et al. Infogan: Interpretable representation learning by information maximizing generative adversarial nets in Proceedings of the 30th International Conference on Neural Information Processing Systems (2016), 2180–2188.

17. Locatello, F. et al. Challenging common assumptions in the unsupervised learning of disentangled representations in international conference on machine learning (2019), 4114–4124.

18. Vinje, W. E. & Gallant, J. L. Sparse coding and decorrelation in primary visual cortex during natural vision. Science 287, 1273–1276 (2000).

19. Perez-Orive, J. et al. Oscillations and Sparsening of Odor Representations in the Mushroom Body. Science 297, 359–365 (2002).

20. Olshausen, B. A. & Field, D. J. Sparse coding of sensory inputs. Current Opinion in Neuro-biology 14, 481–487 (2004).

21. Lewicki, M. S. Efficient coding of natural sounds. Nature Neuroscience 5 (2002).

22. Smith, E. C. & Lewicki, M. S. Efficient auditory coding. Nature 439, 978–982 (2006).

23. Yang, G. R., Cole, M. W. & Rajan, K. How to study the neural mechanisms of multiple tasks. Current opinion in behavioral sciences 29, 134–143 (2019).

24. Yang, G. R., Joglekar, M. R., Song, H. F., Newsome, W. T. & Wang, X.-J. Task representations in neural networks trained to perform many cognitive tasks. Nature neuroscience 22, 297–306 (2019).

25. Caruana, R. Multitask learning. Machine learning 28, 41–75 (1997).

26. Crawshaw, M. Multi-task learning with deep neural networks: A survey. arXiv preprint 2009.09796 (2020).

27. Huang, W., Mordatch, I., Abbeel, P. & Pathak, D. Generalization in Dexterous Manipulation via Geometry-Aware Multi-Task Learning. arXiv preprint 2111.03062 (2021).

28. Van Steenkiste, S., Locatello, F., Schmidhuber, J. & Bachem, O. Are disentangled representations helpful for abstract visual reasoning? arXiv preprint 1905.12506 (2019).

29. Kim, H. & Mnih, A. Disentangling by factorising in International Conference on Machine Learning (2018), 2649–2658.

30. Freedman, D. J. & Assad, J. A. Experience-dependent representation of visual categories in parietal cortex. Nature 443, 85 (2006).

31. Swaminathan, S. K. & Freedman, D. J. Preferential encoding of visual categories in parietal cortex compared with prefrontal cortex. Nature neuroscience 15, 315–320 (2012).

32. Higgins, I. et al. beta-vae: Learning basic visual concepts with a constrained variational framework (2016).

33. Aubry, M., Maturana, D., Efros, A. A., Russell, B. C. & Sivic, J. Seeing 3d chairs: exemplar part-based 2d-3d alignment using a large dataset of cad models in Proceedings of the IEEE conference on computer vision and pattern recognition (2014), 3762–3769.

34. Matthey, L., Higgins, I., Hassabis, D. & Lerchner, A. dSprites: Disentanglement testing Sprites dataset https://github.com/deepmind/dsprites-dataset/. 2017.

35. Yamins, D. L. et al. Performance-optimized hierarchical models predict neural responses in higher visual cortex. Proceedings of the National Academy of Sciences 111, 8619–8624 (2014).

36. Yamins, D. L. & DiCarlo, J. J. Using goal-driven deep learning models to understand sensory cortex. Nature neuroscience 19, 356–365 (2016).

37. Richards, B. A. et al. A deep learning framework for neuroscience. Nature neuroscience 22, 1761–1770 (2019).

38. Lillicrap, T. P. et al. Continuous control with deep reinforcement learning. arXiv preprint 1509.02971 (2015).

39. Raposo, D., Kaufman, M. T. & Churchland, A. K. A category-free neural population supports evolving demands during decision-making. Nature Neuroscience 17, 1784–1792. issn: 1097-6256. NIHMS150003. http://www.pubmedcentral.nih.gov/articlerender.fcgi?artid=4294797%7B%5C&%7Dtool=pmcentrez%7B%5C&%7Drendertype=abstract (2014).

## References

1. Abadi, M. et al. Tensorflow: A system for large-scale machine learning in 12th {USENIX} symposium on operating systems design and implementation ({OSDI} 16) (2016), 265–283.

2. Harris, C. R. et al. Array programming with NumPy. Nature 585, 357–362. https://doi.org/10.1038/s41586-020-2649-2 (Sept. 2020).

3. Pedregosa, F. et al. Scikit-learn: Machine Learning in Python. Journal of Machine Learning Research 12, 2825–2830 (2011).

4. Gao, P. et al. A theory of multineuronal dimensionality, dynamics and measurement. BioRxiv, 214262 (2017).

5. Vinje, W. E. & Gallant, J. L. Sparse coding and decorrelation in primary visual cortex during natural vision. Science 287, 1273–1276 (2000).

6. Perez-Orive, J. et al. Oscillations and Sparsening of Odor Representations in the Mushroom Body. Science 297, 359–365 (2002).

7. Olshausen, B. A. & Field, D. J. Sparse coding of sensory inputs. Current Opinion in Neuro-biology 14, 481–487 (2004).

8. Lewicki, M. S. Efficient coding of natural sounds. Nature Neuroscience 5 (2002).

9. Smith, E. C. & Lewicki, M. S. Efficient auditory coding. Nature 439, 978–982 (2006).

10. Matthey, L., Higgins, I., Hassabis, D. & Lerchner, A. dSprites: Disentanglement testing Sprites dataset https://github.com/deepmind/dsprites-dataset/. 2017.

11. Aubry, M., Maturana, D., Efros, A. A., Russell, B. C. & Sivic, J. Seeing 3d chairs: exemplar part-based 2d-3d alignment using a large dataset of cad models in Proceedings of the IEEE conference on computer vision and pattern recognition (2014), 3762–3769.

12. Lillicrap, T. P. et al. Continuous control with deep reinforcement learning. arXiv preprint 1509.02971 (2015).

13. Higgins, I. et al. β-VAE: Learning basic visual concepts with a constrained variational frame-work in ICLR (2017).

## References

1. Higgins, I. et al. β-VAE: Learning basic visual concepts with a constrained variational frame-work in ICLR (2017).

2. Burgess, C. P. et al. Understanding disentangling in β-VAE. arXiv preprint 1804.03599 (2018).

3. Kim, H. & Mnih, A. Disentangling by factorising in International Conference on Machine Learning (2018), 2649–2658.

4. Locatello, F. et al. Challenging common assumptions in the unsupervised learning of disentan-gled representations in international conference on machine learning (2019), 4114–4124.

5. Higgins, I. et al. beta-vae: Learning basic visual concepts with a constrained variational frame-work (2016).

6. Steegen, S., Tuerlinckx, F., Gelman, A. & Vanpaemel, W. Increasing transparency through a multiverse analysis. Perspectives on Psychological Science 11, 702–712 (2016).

